# Role of the nucleus accumbens in signaled avoidance actions

**DOI:** 10.1101/2024.06.01.596968

**Authors:** Ji Zhou, Sebastian Hormigo, Muhammad S. Sajid, Manuel A. Castro-Alamancos

## Abstract

Animals, humans included, navigate their environments guided by sensory cues, responding adaptively to potential dangers and rewards. Avoidance behaviors serve as adaptive strategies in the face of signaled threats, but the neural mechanisms orchestrating these behaviors remain elusive. Current circuit models of avoidance behaviors indicate that the nucleus accumbens (NAc) in the ventral striatum plays a key role in signaled avoidance behaviors, but the nature of this engagement is unclear. Evolving perspectives propose the NAc as a pivotal hub for action selection, integrating cognitive and affective information to heighten the efficiency of both appetitive and aversive motivated behaviors. To unravel the engagement of the NAc during active and passive avoidance, we used calcium imaging fiber photometry and single-unit recordings to examine NAc GABAergic neuron activity in freely moving mice performing avoidance behaviors. We then probed the functional significance of NAc neurons using optogenetics, and genetically targeted or electrolytic lesions. We found that NAc neurons code contraversive orienting movements and avoidance actions. Intriguingly, specific optogenetic patterns intended to excite NAc GABAergic neurons resulted in local somatic inhibition through GABAergic synaptic collaterals. Nevertheless, these patterns directly excited NAc GABAergic output axons, which in turn inhibited their targets, disrupting active avoidance behavior. Thus, this disruption stemmed from abnormal alterations in the activity of downstream midbrain areas crucial for the behavior. In contrast, direct optogenetic inhibition or lesions of NAc neurons did not impair active or passive avoidance behaviors, challenging the notion of their purported pivotal role in adaptive avoidance. The findings emphasize that NAc is not required for avoidance behaviors, but disruptions in NAc output during pathological states can impair these behaviors.

## Introduction

Animals are often guided by sensory cues to engage in purposeful movements motivated by various contingencies. The execution of avoidance behaviors in response to specific signals represents an adaptive strategy employed in situations where impending danger is indicated. For example, pedestrians navigate a crosswalk when prompted by signals, thereby avoiding potential harm from oncoming traffic. Despite the inherent risk associated with such avoidance behavior, individuals typically execute it routinely, characterized by a sense of caution rather than fear, owing to the perceived control and predictability of the situation (Kamin et al., 1963; Mineka, 1979; Zhou et al., 2022). The exploration of adaptive active avoidance behavior has a rich history spanning nearly a century (Mowrer and Lamoreaux, 1942). However, despite decades of research, a consensus on the neural mechanisms underlying adaptive avoidance remains elusive.

Models outlining the circuitry of active avoidance draw inspiration from the Pavlovian fear conditioning circuit (Davis, 1997; LeDoux, 2000; Maren, 2001; Fanselow and Gale, 2003; Kim and Jung, 2006). In this framework, the conditioned stimulus (CS) traverses from the thalamus to the lateral nucleus of the amygdala, where it forms an association with the aversive stimulus. Subsequently, the signal progresses to the central nucleus, inducing conditioned responses like freezing and potentiated startle. During active avoidance scenarios, a modification of this circuitry is proposed. The CS travels from the lateral nucleus of the amygdala to the basal nucleus and onward to the nucleus accumbens (NAc) under the regulatory influence of the prefrontal cortex (Amorapanth et al., 2000; Moscarello and LeDoux, 2013; Bravo-Rivera et al., 2014; Ramirez et al., 2015). Other theoretical frameworks consider various interacting systems— activation, inhibition, and fight-flight-freeze—as coordinators of avoidance responses (Konorski, 1967; Dickinson et al., 1980; Gray, 2000; Dickinson and Balleine, 2002; McNaughton and Corr, 2004). In this perspective, the NAc is identified as a component of the activating system that drives active responses. In either framework, the NAc assumes a pivotal role in the orchestration of active avoidance behaviors.

The NAc integrates excitatory signals from key forebrain regions, including the medial prefrontal cortex, basolateral amygdala, hippocampus, and thalamus. This intricate interplay is modulated by dopaminergic and GABAergic inputs from the ventral tegmental area (Gerfen and Surmeier, 2011). Predominantly, NAc is composed of GABAergic medium spiny output neurons, while a small fraction (∼5%) of GABAergic interneurons, some releasing acetylcholine, contribute to the neural dynamics within the NAc (Castro and Bruchas, 2019). Various neurochemical and receptor markers delineate the NAc into core and shell regions, each comprising patch and matrix areas housing both direct and indirect pathway medium spiny neurons (Gerfen and Surmeier, 2011; Castro and Bruchas, 2019). The direct pathway influences the substantia nigra and ventral tegmental area in the midbrain, whereas the indirect pathway targets the pallidum and hypothalamus, which subsequently project to the midbrain. These cell groups have been intensely studied revealing distinct features but activate concurrently during behavior (Cui et al., 2013; Klaus et al., 2019). Intriguingly, part of the direct pathway reaches an area in the midbrain pedunculopontine tegmentum (PPT) that is critical for active avoidance, and excitation of these direct pathway GABAergic fibers inhibits PPT and blunts active avoidance (Hormigo et al., 2019), supporting the notion that NAc may play an important role in active avoidance.

Traditionally, the function of the NAc has been closely associated with its dopamine innervation, acting as a mediator of the hedonic aspects of the reward system that propels approach responses. However, this conceptualization has faced challenges, giving way to an alternative framework where the NAc assumes a central role in action selection, integrating cognitive and affective information from various afferent regions. In doing so, it enhances the efficiency and vigor of both appetitive and aversive motivated behaviors (Berridge, 2007; Nicola, 2007; Yin et al., 2008; Salamone and Correa, 2012; Saunders and Robinson, 2012; Floresco, 2015). From the perspective of appetitive, approach, or reward-seeking behavior, the NAc is recognized as a crucial component of an activation and effort-related motivational circuit, influenced by dopamine (Mogenson et al., 1980; Salamone and Correa, 2023). However, a notable gap remains in understanding its purported role during avoidance behaviors, where active or passive responses are motivated by the anticipation of harmful and punishing consequences.

To test the engagement of the NAc during active and passive avoidance behaviors, we measured NAc neuron activity in freely moving mice using calcium imaging fiber photometry, and single-unit recordings. Additionally, we utilized optogenetics and electrolytic, and genetically targeted lesions to probe the functional significance of these neurons during avoidance behaviors. Our findings reveal that NAc neurons code various aspects of signaled avoidance behaviors but are not required to generate these behaviors.

## Results

### NAc neurons code movement direction

Since striatal medium spiny output neurons are GABAergic and activate concurrently during behavior, we targeted them as a coherent population using Vgat-Cre mice (Vong et al., 2011). To assess the population activity of NAc GABAergic neurons, we expressed GCaMP7f (Chen et al., 2013) in these neurons by locally injecting a Cre-AAV in Vgat-Cre mice (n=7). After implanting a single optical fiber within the NAc, we employed calcium imaging fiber photometry, as previously described (Hormigo et al., 2021b; Zhou et al., 2023). Figure 1A illustrates a representative trajectory of the optical fiber, targeting GCaMP-expressing NAc GABAergic neurons. The estimated imaged volume extends ∼200 µm from the optical fiber ending, encompassing ∼2.5×10^7^ µm^3^ (Pisanello et al., 2019).

**Figure 1.**
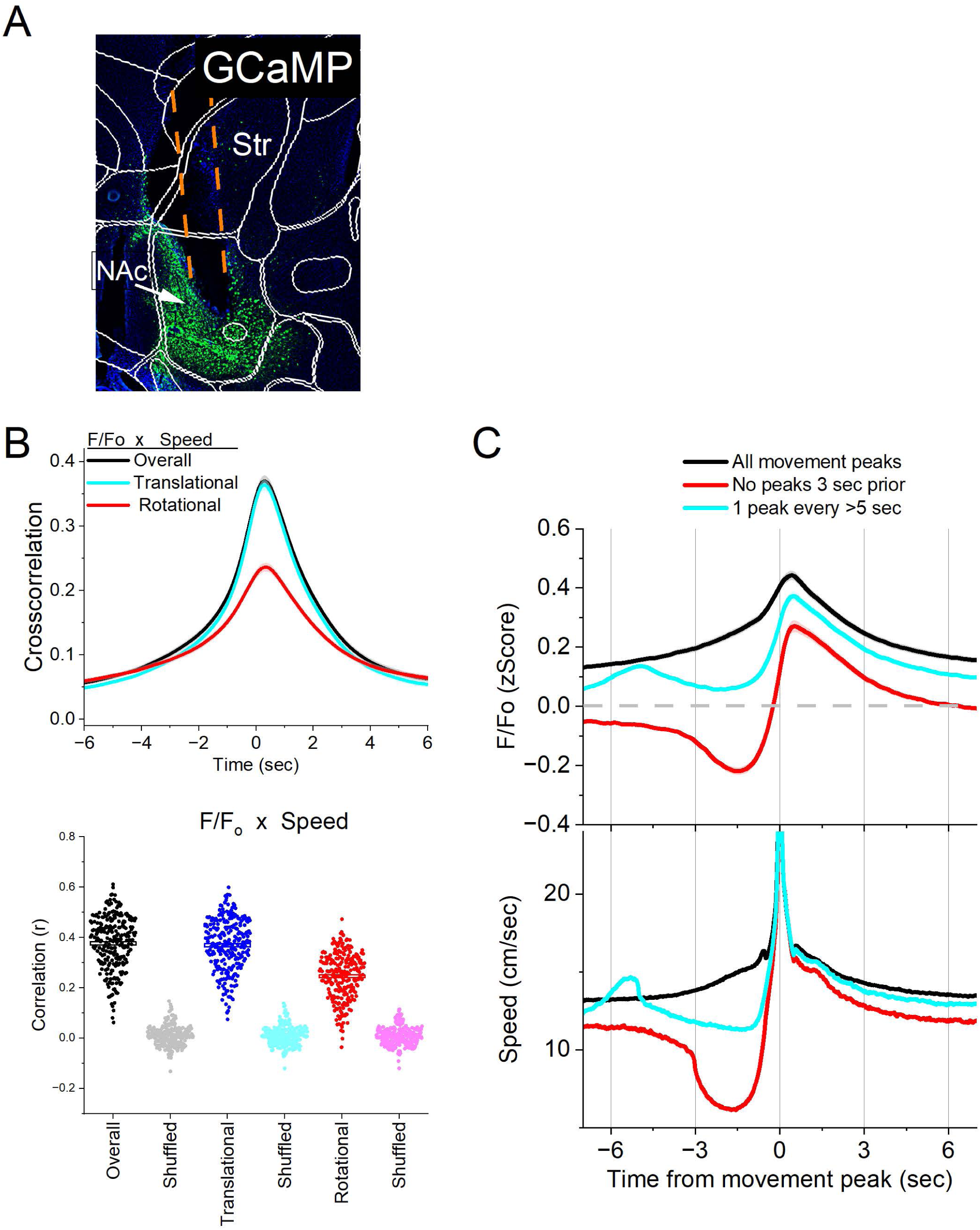
Calcium imaging fiber photometry reveals that NAc GABAergic neurons activate during spontaneous exploratory movement. ***A***, Parasagittal section showing the optical fiber tract reaching NAc and GCaMP7f fluorescence expressed in GABAergic neurons around the fiber ending. The sections were aligned with the Allen brain atlas. **B**, Cross-correlation between movement and NAc F/F_o_ for the overall (black traces), rotational (red) and translational (blue) components (upper panel). Per session (dots) and mean±SEM (rectangle) linear fit (correlation, r) between overall movement and NAc F/F_o_, including the rotational and translational components (lower panel). The lighter dots show the linear fits after scrambling one of the variables (lower panel, shuffled). ***C***, F/F_o_ calcium imaging time extracted around detected spontaneous movements. Time zero represents the peak of the movement. The upper traces show F/F_o_ mean±SEM of all movement peaks (black), those that had no detected peaks 3 sec prior (red), and peaks taken at a fixed interval >5 sec (cyan). The lower traces show the corresponding movement speed for the selected peaks.

In freely moving mice, we conducted continuous measurements of calcium fluorescence (F/F_o_) and spontaneous movement while mice explored an arena. To establish the relationship between movement and NAc activation, cross-correlations were computed between the continuous variables (Fig. 1B upper). Notably, overall movement was strongly correlated with NAc neuron activation. When the movement was dissociated into rotational and translational components, the cross-correlation predominantly involved the translational movement. A linear fit between the movement and F/F_o_ (integrating over a 200 ms window), revealed a strong linear positive correlation that was stronger for the translational component (Fig. 1B lower). These relationships were absent when one variable in the pair was shuffled (Fig. 1B, lower).

To further evaluate the relationship between movement and NAc activation, we detected spontaneous movements and time extracted the continuous variables around the detected movements peaks (Fig. 1C). The detected movements were classified in three categories. The first category includes all peaks (Fig. 1C black traces), which revealed a strong NAc neuron activation in relation to movement. The second category includes movements that had no detected peaks 3 sec prior (Fig. 1C red traces), which essentially extracts movement onsets from immobility. This revealed a sharp activation in association with movement onset. The third category sampled the peaks by averaging every >5 seconds to eliminate from the average the effect of closely occurring peaks (Fig. 1C cyan traces). This category includes movement increases from ongoing baseline movement (instead of movement onsets from immobility) and showed a strong activation of NAc neurons. For the three categories of movement peaks, the NAc activation around movement was significant compared to baseline activity (Tukey p<0.0001). Thus, NAc neurons discharge in relation to the occurrence of movement.

We next determined if the NAc activation during movement depends on the direction of the head movement in the ipsiversive or contraversive direction. Figure 2A shows movement turns in the contraversive (Fig. 2A cyan) and ipsiversive (Fig 2A red) directions versus the recorded NAc neurons. While the detected turns were opposite in direction and similar in amplitude, the NAc neuron activation was sharper when the head turned in the contraversive direction. We compared the area, peak amplitude, and peak timing of the F/Fo activation between ipsiversive and contraversive turns (Fig. 2B). For *all turns, no turns 3 sec prior, and 1 turn per >5 sec*, the F/Fo area (Tukey t(256)= 6.16 p<0.0001, Tukey t(256)= 4.99 p=0.0005, and Tukey t(256)= 5.22 p=0.0003) and amplitude (Tukey t(256)= 6.82 p<0.0001, Tukey t(256)= 5.89 p<0.0001, and Tukey t(256)= 6.4 p<0.0001) of contraversive turns were larger than ipsiversive turns. However, the times to peak did not differ. Therefore, NAc GABAergic neurons code movement direction discharging more sharply to contraversive turns.

**Figure 2.**
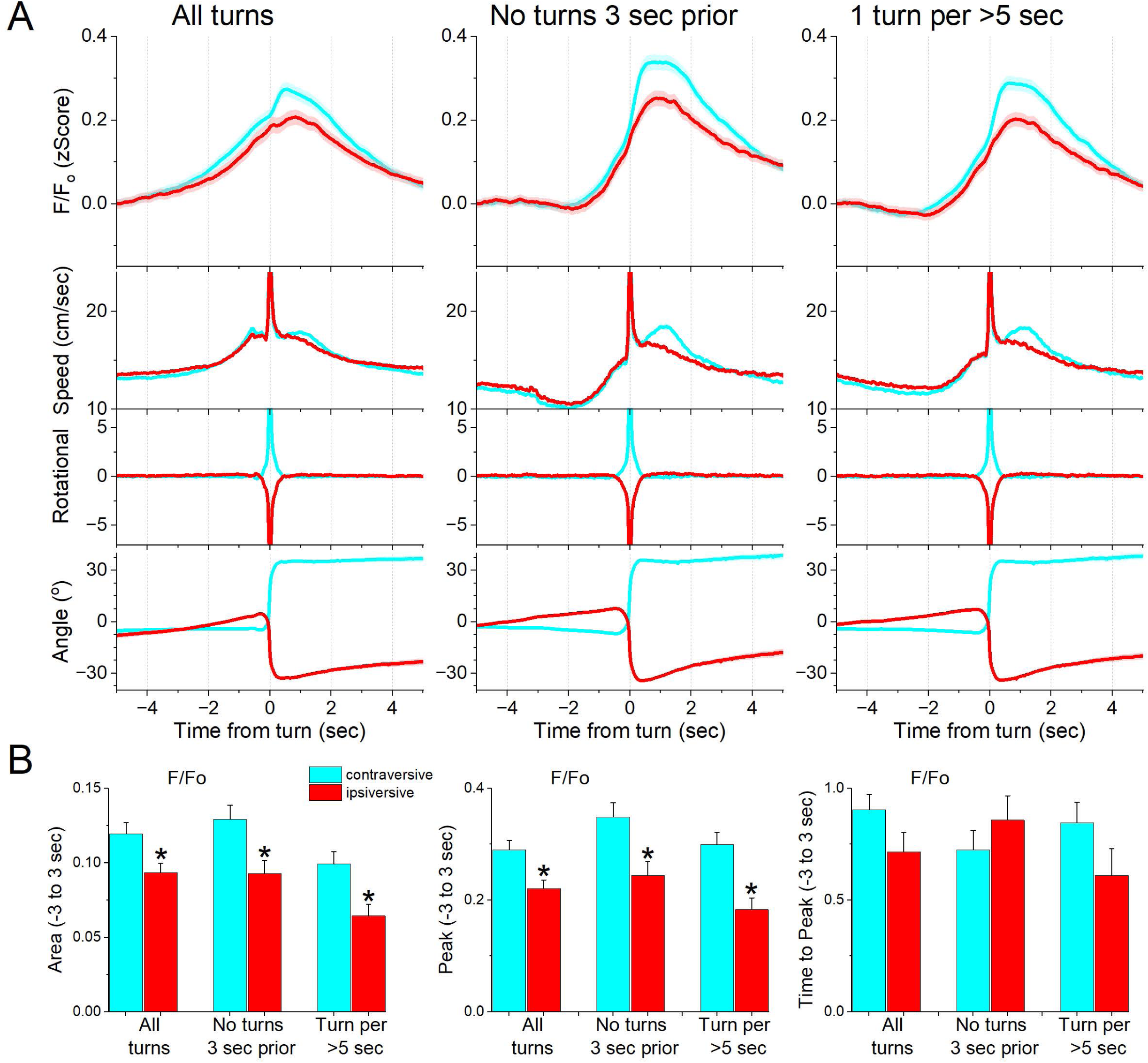
NAc GABAergic neurons code the direction of spontaneous contraversive exploratory turning movements. ***A***, F/F_o_ calcium imaging, overall movement, rotational movement, and angle of turning direction for detected movements classified by the turning direction (ipsiversive and contraversive; red and cyan) versus the side of the recording (implanted optical fiber). At time zero, the animals spontaneously turn their head in the indicated direction. The columns show all turns (left), those that included no turn peaks 3 sec prior (middle), and peaks selected at a fixed interval >5 sec (right). Note that the speed of the movements was similar in both directions. ***B***, Population measures (area of traces 3 sec around the detected peaks) of F/F_o_ and movement (overall, rotational, and translational) for the different classified peaks. The measures that were different (p<0.05) between ipsiversive and contraversive movements are indicated by *.

### NAc neurons activate during goal-directed avoidance movement

The results indicate that NAc neurons activate during movement. Therefore, we explored the activation of NAc neurons during a series of cued (signaled) avoidance procedures (AA1, AA2, AA3, and AA4 (Zhou et al., 2022; Zhou et al., 2023)) in which mice move (actively avoid) or postpone movement (passively avoid) to prevent an aversive US. Since these procedures are signaled by tones, we first conducted auditory mapping sessions to test if auditory tones activated NAc neurons in freely behaving mice.

During auditory mapping sessions (42 sessions in 7 mice; Fig. 3A,B), mice were placed in a small cage (half the size of a shuttle box) and 10 auditory tones of different saliencies, defined by sound pressure level (SPL in dB; low and high, ∼70 and ∼85 dB) and frequency (4, 6, 8, 12, 16 kHz), were presented in random order (1 sec tones every 4-5 sec, each repeated 10 times per session). The calcium signal evoked in NAc neurons by the tones (0-1.5 sec window) depended on SPL (2WayRMAnova F(1,41)= 61.33 p<0.0001) and frequency (F(4,164)= 10.05 p<0.0001), and these factors interacted (F(4,164)= 5.53 p=0.0003). In general, higher SPL and frequency tones produced stronger NAc activation (Fig. 3), but the lowest frequency (4 kHz) had a weaker activating effect that was not as sensitive to SPL as the higher frequencies (Tukey t(164)=1.9 p=0.2 Lo vs Hi at 4 kHz). However, these same effects were observed on movement (overall speed). Thus, the movement evoked by the tones also depended on tone SPL (F(1,41)= 76.81 p<0.0001) and frequency (F(4,164)= 31.79 p<0.0001), and these factors interacted (F(4,164)= 3.94 p=0.0044). The movement evoked by the tones consisted of both rotational and translational components (Fig. 3B), and each of these components showed the same effects as overall movement (SPL or frequency p<0.0001). However, it is worth noting that the amplitudes of F/Fo activation (0.1-0.2 zScore) and movements (∼2-4 cm/sec) evoked by the tones are relatively small. The results indicate that NAc neurons respond to salient auditory tones in association with movement.

**Figure 3.**
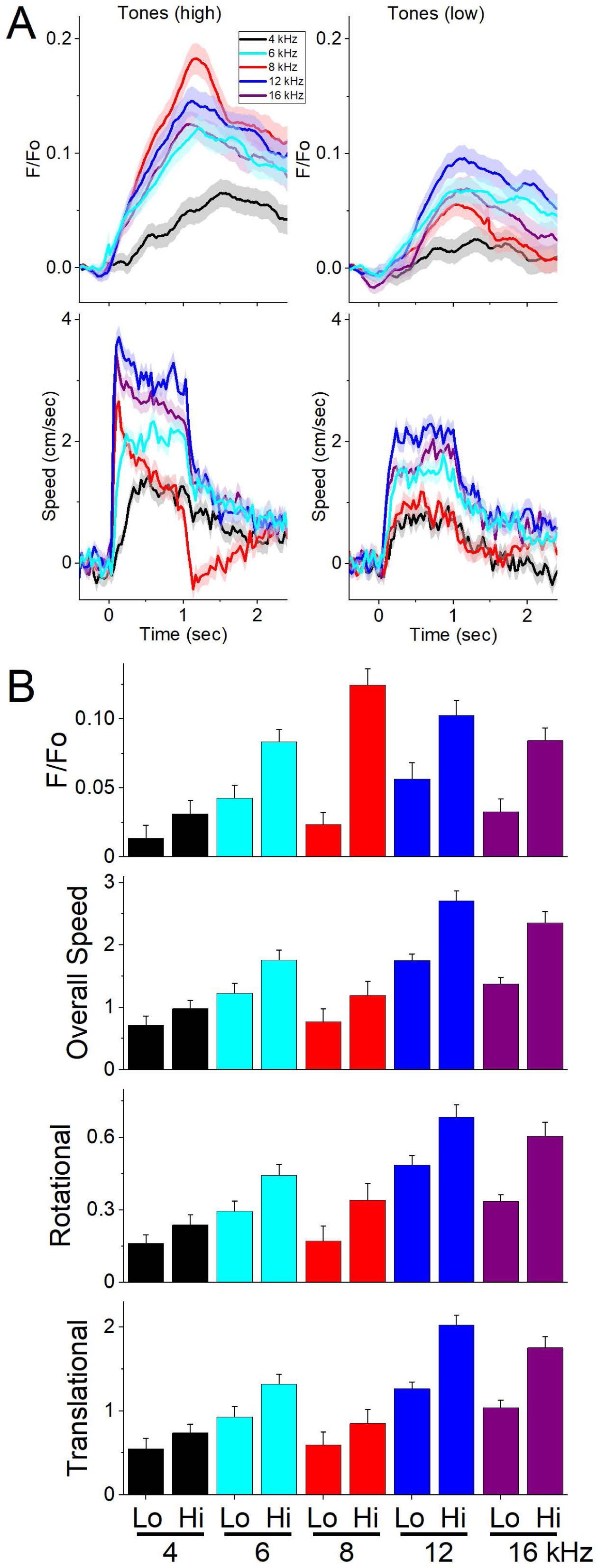
NAc GABAergic neurons discharge to auditory tones in association with movement. ***A***, Example F/F_o_ calcium imaging and movement traces (mean±SEM) evoked from NAc neurons by auditory tones (1 sec) of different saliency. The tones vary in frequency (4-16 kHz) and SPL (low and high dB). ***B***, Area of F/F_o_, overall movement, and movement components (rotational and translational) measured during a time window (0-2 s) after tone onset.

We then measured NAc neuron activation as the mice performed the avoidance procedures in a shuttle box (Fig. 4A). Figure 4B shows the behavioral performance of the animals in these tasks including the percentage of avoids (black circles), the avoid latencies (orange triangles), and the number of intertrial crossings (cyan bars). As previously shown, animals perform a large percentage of active avoids during the AA1, AA2, AA3-CS1, and AA4 procedures (Hormigo et al., 2021b). During AA2, ITCs are punished. Consequently, avoid latencies shift longer in an apparent reflection of caution (Zhou et al., 2022). During AA3, animals learn to discriminate between two CS’s, passively avoiding when AA3-CS2 is presented. During AA4, the avoid latencies adapt to the varying duration of three active avoid intervals (4, 7, and 15 sec) signaled by different CS tones.

**Figure 4.**
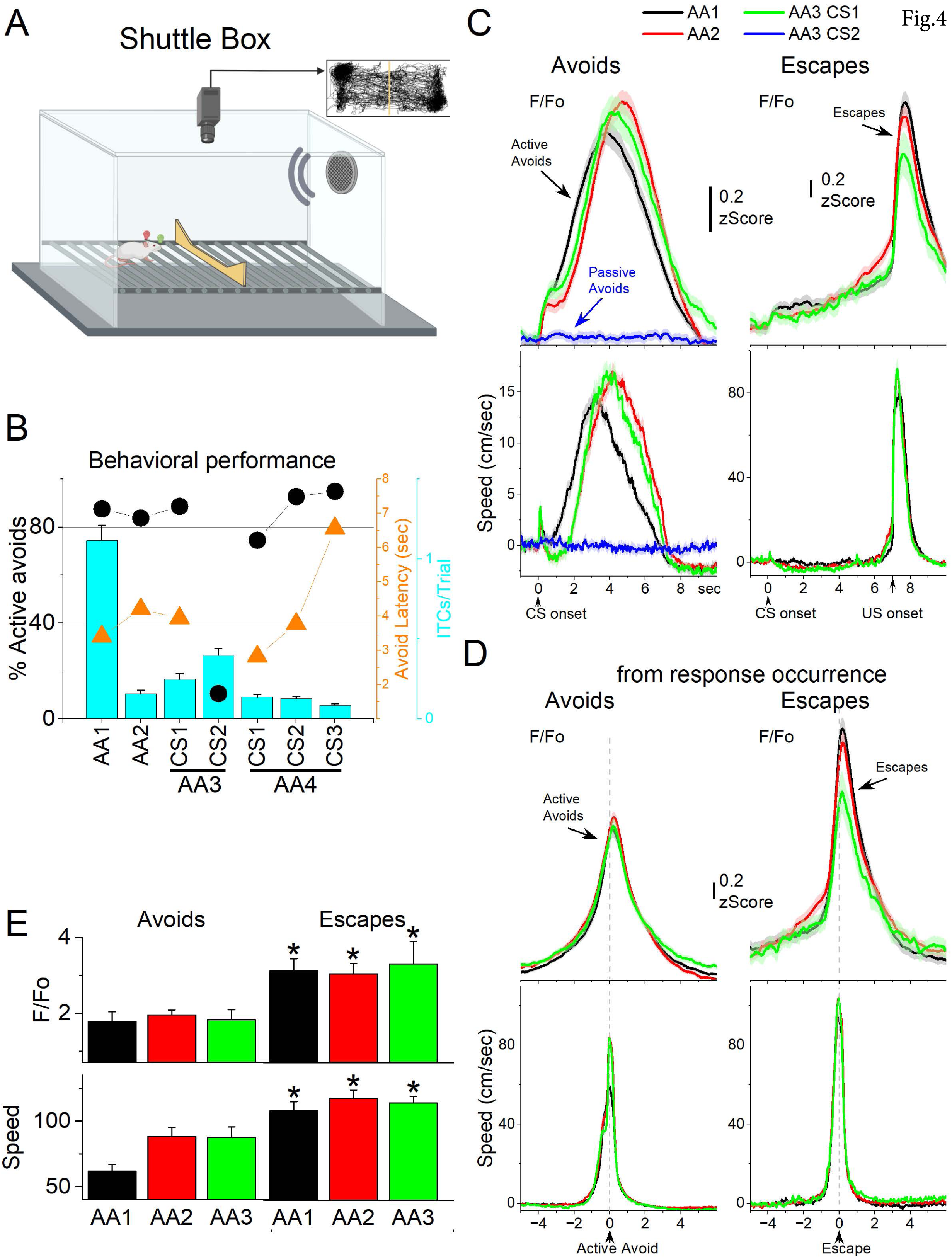
NAc GABAergic activation in the context of signaled active avoidance. ***A***, Arrangement of the shuttle box used during signaled avoidance tasks. ***B***, Behavioral performance during the four different avoidance procedures showing the percentage of active avoids (black circles), avoidance latency (orange triangles), and ITCs (cyan bars). ***C***, F/F_o_ and overall movement traces from CS onset for AA1, AA2 and AA3 (CS1 and CS2) procedures per trials classified as avoids (left) or escapes (right) of CS-evoked orienting responses measured by tracking overall head speed. ***D,*** Same as C from response occurrence. ***E***, Population measures of F/Fo and speed for avoids and escapes during AA1, AA2 and AA3 (CS1). Asterisks denote p<0.05 vs Avoids.

Figure 4C shows F/Fo and movement traces from CS onset for the AA1 (black), AA2 (red), and AA3 (green) avoidance procedures classified as correct responses (left panel, avoids; for AA3-CS2 correct passive avoids are shown in blue) or errors (right panel, escapes). During the avoidance procedures, the CS that drives active avoids (Fig. 4C black, red, green) caused a sharp and fast (<0.5 sec) F/Fo peak at CS onset, which is associated with the typical orienting head movement evoked by the CS depending on task contingencies and SPL (Zhou et al., 2023). AA3-CS2, which drives passive avoids, produced nil NAc activation at CS2 onset in association with nil orienting head movement due to lower SPL. Thus, the CS onset NAc activation evoked by CS2 was smaller than the activation evoked by CS1 (Tukey t(14)= 4.53 p=0.0064).

The succeeding avoidance movement was associated with strong NAc neuron activation (Fig. 4C avoids). In AA1, there was a large F/Fo peak related to the avoidance movement. As mice transitioned to AA2, the NAc activation shifted to the right following the delayed avoid latencies characteristic of this procedure (Fig. 4C black vs red traces). Thus, the activation of NAc neurons is closely associated with the active avoid movement. In addition, when mice failed to avoid, NAc neurons activated very sharply in association with the fast escape responses evoked by the US during the escape interval (Fig. 4C escapes). Furthermore, during the AA3 procedure, only CS1, which drives active avoids, produced strong NAc activation (Fig. 4C green). CS2, which drives passive avoids, produced nil NAc activation (Fig. 4C blue). To measure avoid and escape responses, we extracted the F/Fo and speed from the response occurrence for AA1, AA2, and AA3 (Fig. 4D). Both avoids and escapes were associated with robust NAc activation during the three procedures (Fig. 4D,E). However, since escapes occurred at higher speeds (Tukey t(6)= 12.04 p=0.0001 avoids vs escapes), they were associated with stronger NAc activation than avoids (Tukey t(6)= 6.65 p=0.0033). Moreover, as is typical (Zhou et al., 2022), the speed of avoids was higher during AA2 compared to AA1 (t(12)= 4.85 p=0.0129 AA1 vs AA2), but this was not consistently associated with a larger NAc activation during AA2.

During AA4, mice adapt their avoidance movement to the duration of the avoidance interval signaled by each of the three CSs (Hormigo et al., 2021b). Accordingly, NAc activation shifted to reflect the avoidance movement (Fig. 5A). When responses were measured from response occurrence (Fig. 5A right panels and 5B), the CS1 avoids speed was faster than CS2 (Tukey t(36)= 5.3 p=0.001) or CS3 (Tukey t(36)= 5.2 p=0.001), consistent with the more imminent threat signaled by CS1, which has a shorter 4-sec avoidance interval. However, this was not reflected in a difference of the peak activation of NAc neurons (RMAnova F(2,36)= 0.72 p=0.48).

**Figure 5.**
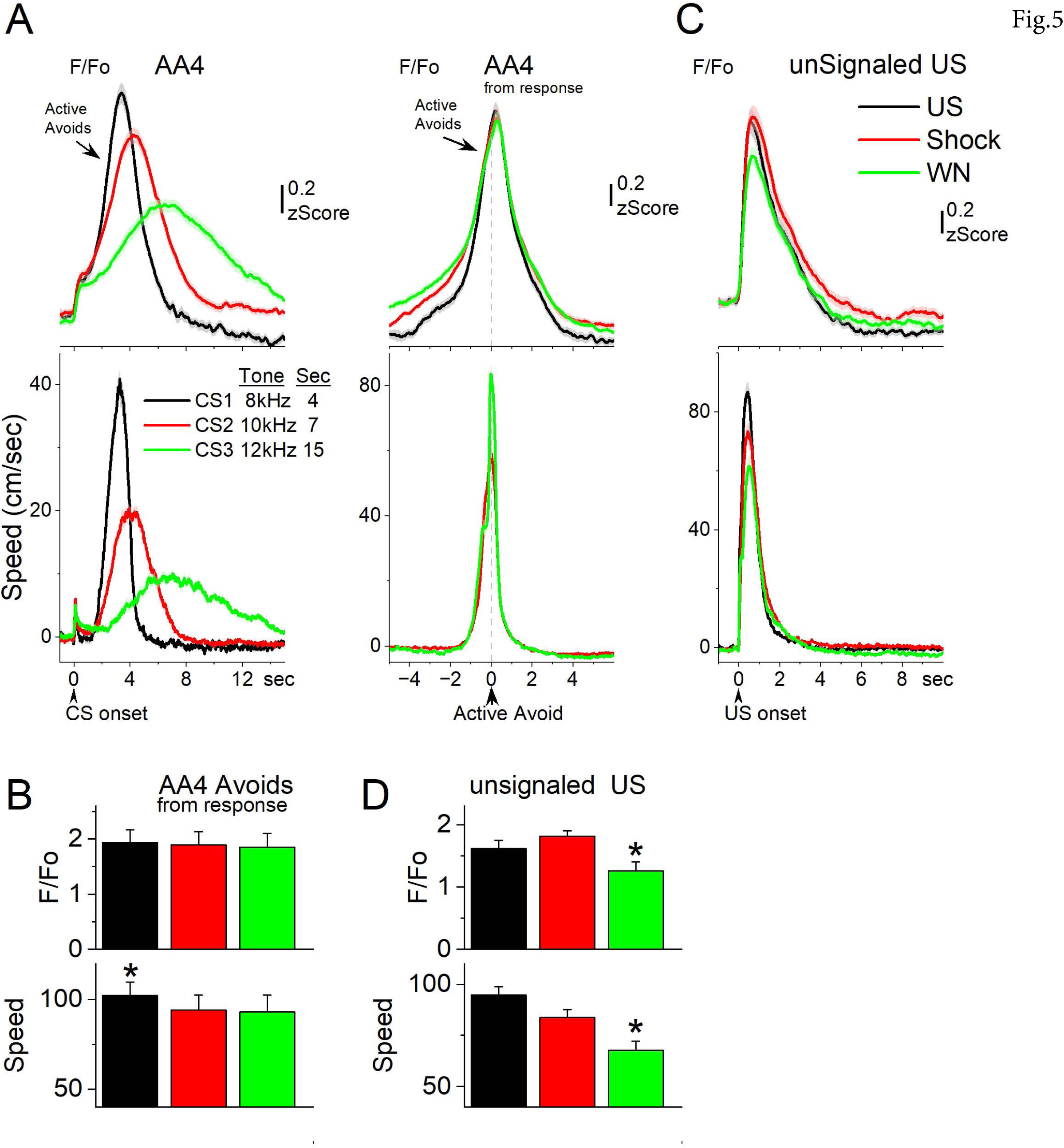
NAc GABAergic neurons track the avoidance and escape movement. ***A***, F/F_o_ and overall movement traces from CS onset (left) and response occurrence (right) for avoids during the AA4 procedure, which include three CSs that signal avoidance intervals of different durations. ***B***, Population measures (–3 to 3 sec area, mean ± SEM) from response occurrence. Asterisks denote significant differences vs other stimuli. ***C***, F/F_o_ and overall movement traces from US onset for escapes during the unsignaled US procedure, which includes the US, or each of its components delivered alone (foot-shock and white noise). ***D***, Population measures (0 to 5 sec area, mean ± SEM) for the data in C. Asterisks denote significant differences vs other CS’s.

Since mice have high rates of avoids in these tasks, and consequently relatively few escapes, we conducted additional sessions (Fig. 5C,D; 7 mice) in which the US (foot-shock and white noise) was presented unsignaled to drive an escape on every trial. To distinguish the contribution of the foot-shock and white noise presented by the US, additional trials in the same session presented the foot-shock or the white noise alone. The unsignaled US evoked a strong NAc activation in association with the fast escapes. Presentation of the foot-shock alone produced NAc activation and escapes like the full US (Fig. 5C,D). However, presentation of the white noise alone drove escapes at lower speeds, which were associated with less NAc activation than the foot-shock alone (Tukey t(32)= 6.6 p=0.00014 speed; Tukey t(32)= 4.8 p=0.004 F/Fo) or the full US (Tukey t(32)= 11.2 p<0.0001 speed; Tukey t(32)= 6.3 p=0.0002 F/Fo). Thus, the NAc activation during unsignaled US presentations follows the speed of the escape movement driven by the stimuli.

In conclusion, NAc neurons activate at CS onset in association with an orienting movement, and then discharge more robustly during the ensuing active avoid and escape movements. NAc activation has the potential to drive active avoids, but NAc activation may instead reflect the ongoing movement.

### NAc electrophysiology support the calcium imaging results

To validate the primary findings from calcium imaging, which indicate that NAc neurons become active during orienting and avoidance movements, we conducted electrophysiological recordings of unit activity in the NAc while mice engaged in signaled avoidance. To ensure accurate placement of the recording electrodes within the NAc, a marking lesion was created following the final recording session (Fig. 6A). We identified a set of multi-unit clusters recorded over multiple sessions for analysis (n=6 recording electrodes from 3 mice).

**Figure 6.**
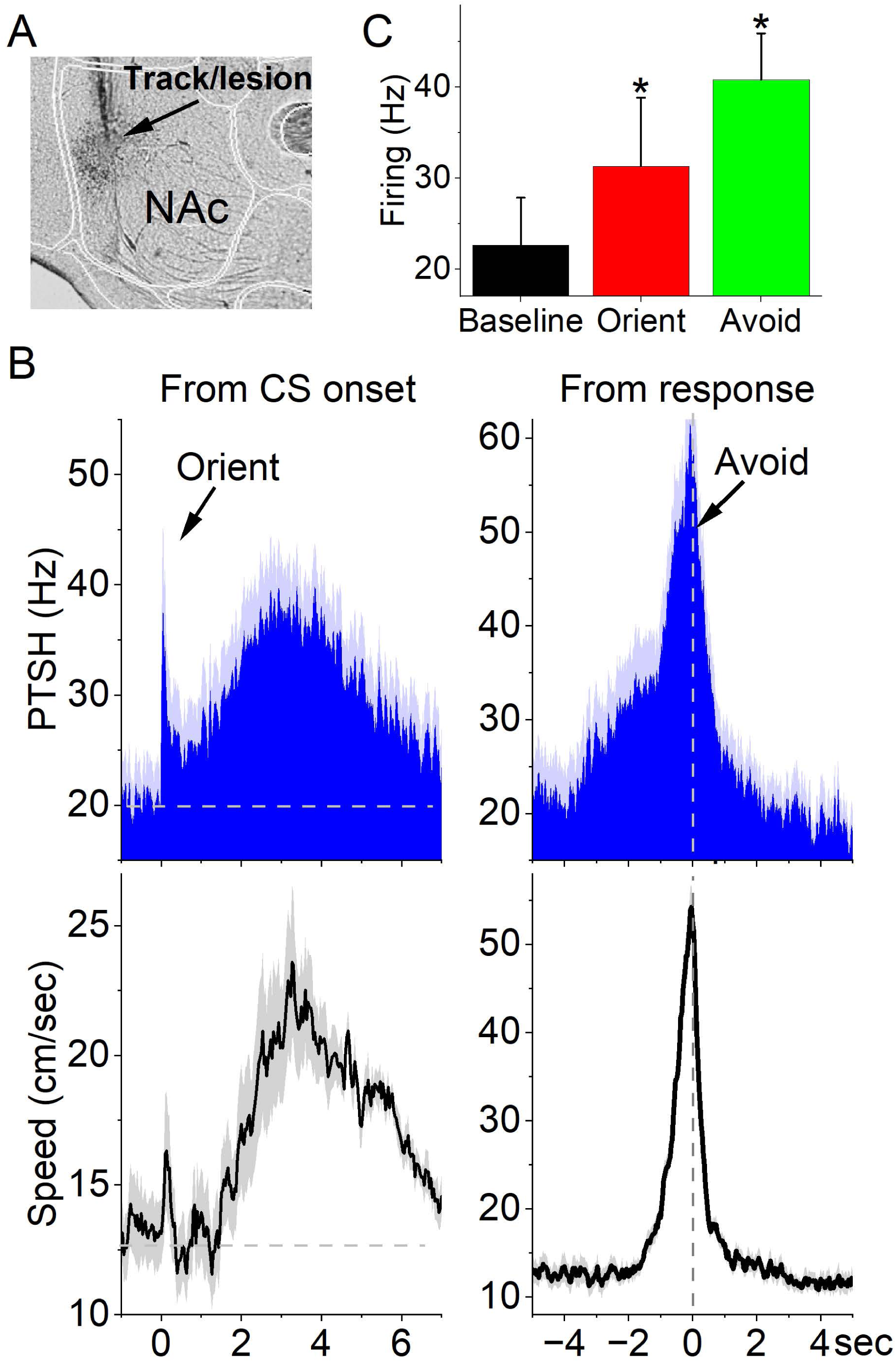
NAc multi-unit activity measured with electrophysiology in freely moving mice performing signaled active avoidance resembles the calcium imaging activity. ***A***, Bright-field sagittal section aligned with the Allen Brain Atlas showing a marking lesion in the NAc in a static NAc recording electrode. ***B***, PSTH of unit firing (Hz) and movement (speed) traces showing modulation during signaled active avoidance (AA1/2). The PSTH is calculated from CS onset or from response occurrence. The lower panels show overall movement. ***C***, Population measures of unit firing (left) during orienting response (0-0.5 sec) and avoidance response (from response –2 to 2 sec) windows compared to baseline firing (pre-CS).

During active avoidance (AA1/2), NAc neurons discharged in a way that resembled the activation revealed by calcium imaging. From CS onset (Fig. 6B,C), there was a transient sharp discharge associated with the orienting head movement (Tukey t(5)= 4.13 p=0.03 baseline vs 0-0.5 sec after CS onset), followed by a subsequent robust activation during the avoidance movement. From response occurrence (Fig. 6B,C), there was a robust activation related to the avoidance response, which resembled the calcium imaging activation (Tukey t(5)= 5.4 p=0.01 baseline vs 2 sec around avoid occurrence). These results support the calcium imaging findings indicating that NAc neurons activate primarily during movement, including orienting and avoidance responses. This aligns with previous work conducted on freely behaving rodents, which has demonstrated that striatal neurons discharge during movement and associated somatosensory input (Carelli and West, 1991; Cui et al., 2013; Markowitz et al., 2018; Robbe, 2018; Sales-Carbonell et al., 2018).

### Excitation of NAc GABAergic neurons blocks signaled avoidance

The preceding experiments using imaging and electrophysiology recordings indicate that NAc neurons discharge during signaled avoidance and could have an important role in producing avoidance behavior. Therefore, we used optogenetics to modulate (excite or inhibit) NAc GABAergic neurons as mice performed AA1, AA2, and AA3 procedures. First, we tested the effects of green and blue light in No Opsin mice (n=10) subjected to the same behavioral procedures, which revealed little effect on behavioral performance (active avoids rate and latency) in AA1, AA2, or AA3 (RMAnova F(1,9)<3 p>0.15).

To excite NAc GABAergic neurons, we expressed ChR2 in the NAc of Vgat-Cre mice with bilateral injections of a Cre-inducible AAV (Fig. 7A; Vgat-NAc-ChR2, n=10 mice). Excitation of NAc neurons during the avoidance intervals of AA1 (Fig. 7B) decreased the percentage of active avoids when high frequency (>20 Hz trains of 1 ms pulses; Tukey t(72)> 10.4 p<0.0001 CS vs CS+Light at 40, 66 or 100 Hz) or Cont (Tukey t(72)= 8.31 p<0.0001 CS vs CS+Light Cont) blue light was applied. The suppression was maximal during blue light trains at 66 Hz, and sensitive to light power (2WayRMAnova interaction 8 CS+Light patterns x 2 Powers F(7,63)= 5.75 p<0.0001; Fig. 7B Green vs Red symbols). At low power, Cont blue light had no effect on active avoids (Tukey t(72)= 3.08 p=0.44 CS vs CS+Light Cont at Low power) but they were strongly suppressed by 66-100 Hz trains (Tukey t(72)= 6.42 p<0.0001 CS vs CS+Light 66 Hz at Low power). At high powers, Cont blue light suppressed avoids (Tukey t(72)= 5.8 p<0.0001 CS vs CS+Light Cont at High power) but significantly less than 40-100 Hz trains (Tukey t(72)= 11.3 p<0.0001 CS vs CS+Light 66 Hz at High power; Tukey t(72)= 5.44 p=0.0001 Cont vs 66 Hz at High power). Thus, blue light high frequency trains (40-100 Hz) are more effective at suppressing avoids than Cont blue light. As revealed later, this indicates a complex effect of the blue light on NAc firing because Cont is typically the light pattern that drives the strongest firing when applied to the somatodendritic region of ChR2-expressing neurons.

**Figure 7.**
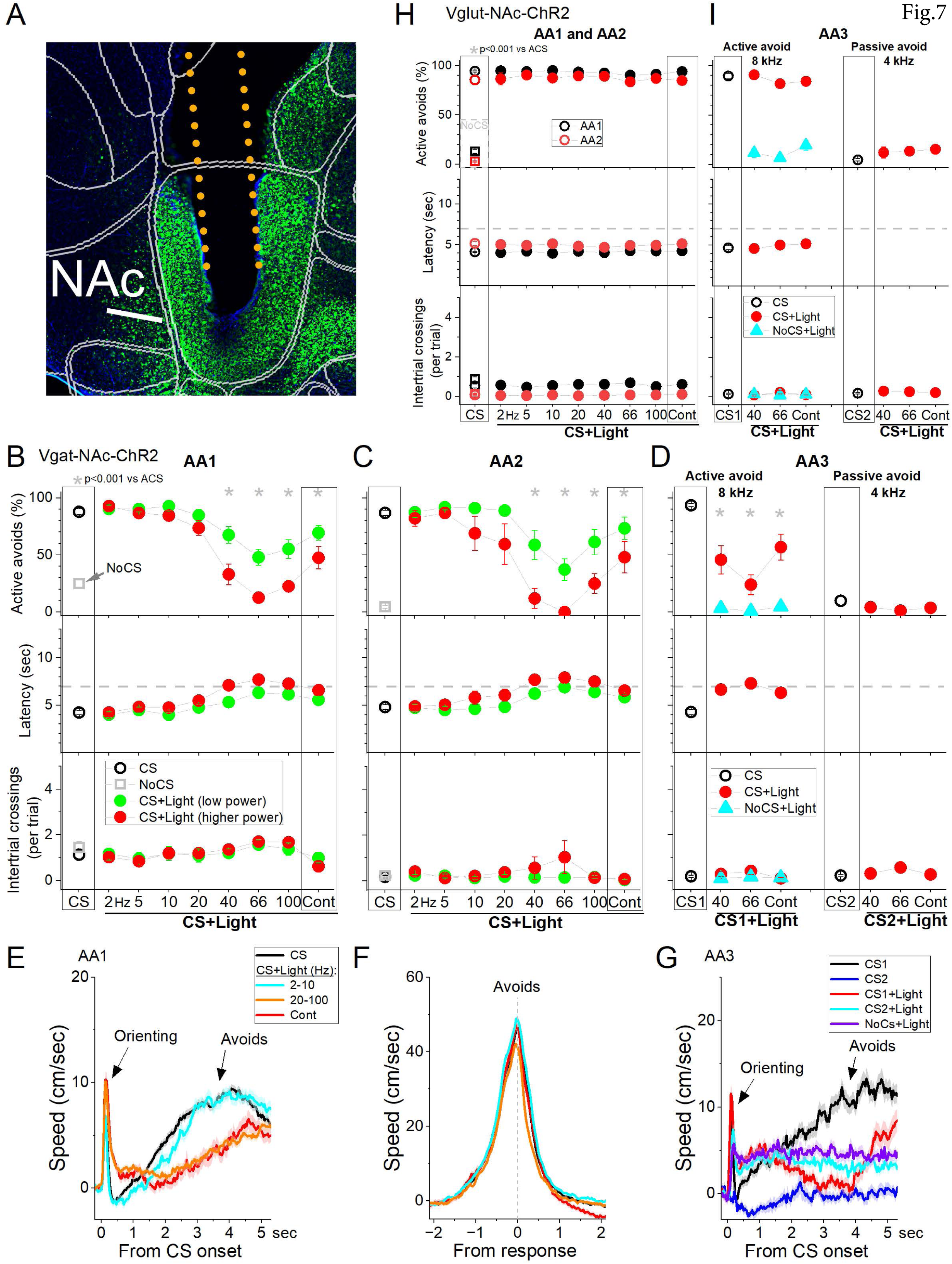
Optogenetic activation of ChR2-expressing NAc GABAergic neurons impairs signaled avoidance. ***A***, Parasagittal section showing the optical fiber tract reaching NAc and ChR2 fluorescence expressed in GABAergic neurons. The section was aligned with the Allen brain atlas. ***B,C,D,*** Effects of patterns of blue light applied in NAc as low (green circles) or higher (red circles) power during AA1 (***B***), AA2(***C***), and AA3 (***D***). CS trials are control trials without light. CS+Light trials include the CS and light patterns (trains of 1-ms pulses at the noted frequencies in Hz or Cont). During AA3, CS1 trials are active avoidance trials (like CS trials in AA1/2) and CS2 trials are passive avoidance trials. The plots show the percentage of active avoids, the response latency, and the intertrial crossings per trial. NoCS trials are catch trials without CS or consequence. ***E,*** Traces of overall speed during AA1 for CS trials, and CS+Light trials combined according to the light pattern into low frequency (2-10 Hz), high frequency (20-100Hz), and Cont. The trials are aligned by CS onset, which reveals the orienting response evoked by the CS and the subsequent action. ***F,*** Same as *E* but traces are aligned by the avoidance response occurrence, which reveals the peak speed of the avoid action. ***G,*** same as *E* but for AA3. CS1+Light and CS2+Light trials show the effect of the Light patterns combined (40-66 Hz and Cont). Note that the light activation of NAc GABAergic neurons increases overall speed in NoCS+Light trials, indicating that it enhances movement whilst impairing active avoidance responding. ***H,I,*** Same as ***B*** and ***D*** but for a control group of mice that do not express ChR2 in NAc as they were injected with an AAV targeting glutamatergic neurons. The light patterns have negligible effects on AA1 (***H***) and AA3 (***I***).

Despite the strong blockade of avoids driven by the CS, the same light patterns continuing during the escape interval (7 sec after CS onset when avoid fails) did not interfere with the escapes driven by the US. Thus, when mice fail to avoid, they escape rapidly (Fig. 7B latency middle panel). As is typical when mice are exposed to the US, the number of ITCs increased slightly during the ITI after CS+Light trials at 66 Hz (Fig. 7B ITCs lower panel; Tukey t(72)= 5.27 p=0.011 CS vs 66 Hz). The suppression of active avoids was similar in AA2 (Fig. 7C) and AA3-CS1 (Fig. 7D). Moreover, signaled passive avoids during AA3-CS2 were not affected by excitation of NAc GABAergic neurons with any of the three patterns tested (Fig. 7D; Tukey t(42)<1.38 p>0.97 CS2 vs 40, 66 Hz or Cont). Finally, application of these same light patterns during AA3 in NoCS+Light trials (i.e., without CS/US) showed no effects (Fig. 7D cyan triangles) on the measured avoidance parameters.

Movement tracking revealed that the effects of NAc excitation on signaled active avoidance (AA1, AA2 and AA3-CS1) were associated with an increase in the orienting response amplitude (Fig. 7E,G; Tukey t(9)= 4.5 p=0.01 CS vs CS+Light >20 Hz and Cont). To measure the peak speed of avoids and escapes we used the response occurrence marker (when mice cross the door; Fig. 7F) because these responses may occur at different times from CS onset. The peak speed of avoids (Tukey t(9)= 0.09 p=0.97) or escapes (Tukey t(9)= 2.12 p=0.17) measured from response occurrence were not affected by NAc excitation when they occurred. This suggests that mice may not have a movement generation impairment during NAc GABAergic neuron excitation. In AA3-CS2 trials, we evaluated the effects of NAc excitation on movement during signaled passive avoids. Intriguingly, NAc GABAergic neuron excitation produced an increase in movement (Fig. 7G, Tukey t(18)= 5.6 p=0.004 AA3-CS2 vs AA3-CS2+Light). However, the increased movement was restricted to the compartment where the animals are located because as already noted the number of errors (i.e., active avoids during AA3-CS2 trials) did not increase. While mice decreased their movement during control AA3-CS2 signaled passive avoidance trials (Fig. 7G blue), the excitation of NAc GABAergic neurons increased movement during both AA3-CS2+Light (Fig. 7G cyan) and NoCS+Light (Fig. 7G purple) trials. Thus, NAc GABAergic excitation increases movement, which is associated with impaired active avoidance, but not passive avoidance.

To determine the selectivity of exciting NAc GABAergic neurons on avoidance, we injected the same AAV in Vglut2-Cre mice (n=2) or a non-Cre AAV with a CaMKII promoter in C57BL/6J mice (n=4), which can target glutamatergic neurons. As expected, these injections did not show significant ChR2 expression within NAc because Vglut2 expression in NAc is nil (Vong et al., 2011), and therefore we formed a combined control group (Vglut-NAc-ChR2, n=6 mice) that underwent the same protocols as the Vgat-NAc-ChR2 mice. Blue light had little effect on active avoids during AA1 (Fig. 7H black, RMAnova F(8,40)= 1.28 p=0.27), AA2 (Fig. 7H red, RMAnova F(8,40)= 0.86 p=0.55), AA3-CS1 (Fig. 7I left, RMAnova F(3,15)= 1.9 p=0.15), and AA3-CS2 (Fig. 7I right, RMAnova F(3,15)= 2.5 p=0.09).

Taken together, these results demonstrate that high frequency trains of blue light applied to NAc GABAergic neurons strongly suppress signaled active avoidance but has little effect on signaled passive avoidance. These effects are analogous to the selective excitation of direct pathway neuron axons that originate in NAc and reach the midbrain, which also suppresses signaled active avoidance (Hormigo et al., 2021c). This suppression of active avoidance is not caused by movement inhibition because mice escape the US without impairment, and blue light trains increase movement. Cont blue light produced similar but weaker effects than trains, which led us to test next how Cont and high frequency trains affect NAc neural activity in Vgat-NAc-ChR2 mice.

### Excitation of NAc GABAergic neurons produces local recurrent inhibition

Previous work has shown that patterns of blue light, such as Cont and trains of 1-ms pulses at different frequencies, have distinct effects on the firing of ChR2-expressing GABAergic networks (Hormigo et al., 2016; Hormigo et al., 2019). When the blue light is applied to the somatodendritic area of ChR2-expressing neurons, Cont enhances the firing rate more strongly than high frequency trains. Conversely, when the blue light is applied to axon fibers of ChR2-expressing GABAergic neurons, high frequency trains (40-66 Hz) are much more effective than Cont at driving sustained IPSPs in the postsynaptic neurons (Hormigo et al., 2019). Consequently, when the blue light patterns target an area that contains both ChR2-expressing GABAergic neurons and fibers (collaterals and/or afferents), Cont and trains reveal their differential effects (Hormigo et al., 2021a). For example, in SNr, the same GABAergic neurons discharge robustly during Cont, due to direct ChR2 activation, but are recurrently inhibited by high frequency trains, due to sustained IPSPs from the synaptic collaterals and afferents that express ChR2 (Hormigo et al., 2021a). We reasoned that this may be occurring in the NAc of Vgat-NAc-ChR2 mice when subjected to Cont and trains of blue light, because NAc consists of networks of GABAergic neurons, some of which have strong intra-NAc synaptic connections (Tepper et al., 2004; Castro and Bruchas, 2019), which provides the substrate for high frequency trains to inhibit NAc principal neurons in Vgat-NAc-ChR2 mice. To determine if this is the case, we performed whole-cell recordings from principal spiny stellate neurons in slices of Vgat-NAc-ChR2 mice and tested the effects of Cont and trains of blue light on PSPs.

Single (1 ms) pulses of blue light produced an intrinsic EPSP, driven by direct activation of ChR2 channels, followed by a sharp synaptic IPSP that curtailed the EPSP (Fig. 8A; see inset for a close-up). Trains of pulses drove these single responses, but as the train frequency increased sustained IPSPs strongly hyperpolarized the cells and suppressed firing, as shown previously (Hormigo et al., 2021a). The suppression was maximal at 40-66 Hz and waned at 100 Hz as the number of pulses increased, which drives more depolarizing current mediated by direct ChR2 activation. Fig. 8B,C show PSPs evoked by Cont and single pulses of blue light at a low and a high power. Notably, the evoked IPSPs are stronger during the single 1-ms pulses because Cont activates more depolarizing ChR2 current directly in the neuron. Fig. 8D compare the effect of Cont and a 40 Hz train (500 ms) on a neuron that is activated to fire by positive current injection (from 200 ms before to 100 ms after the blue light). The 40 Hz train was much more effective than Cont at suppressing firing, although the inhibitory currents driven by Cont also shunted neuronal firing, but much less than the trains. When neurons were in a deactivated state (nil or little spontaneous firing) 40-66 Hz trains had little effect but Cont increased firing (Fig. 6E, n=4 cells, Wilcoxon p=0.03). However, when neurons were activated by current injection, trains at 40-66 Hz sharply inhibited firing (p=0.01 Spont vs 40-66 Hz), more effectively than Cont (p=0.04 Spont vs Cont).

**Figure 8.**
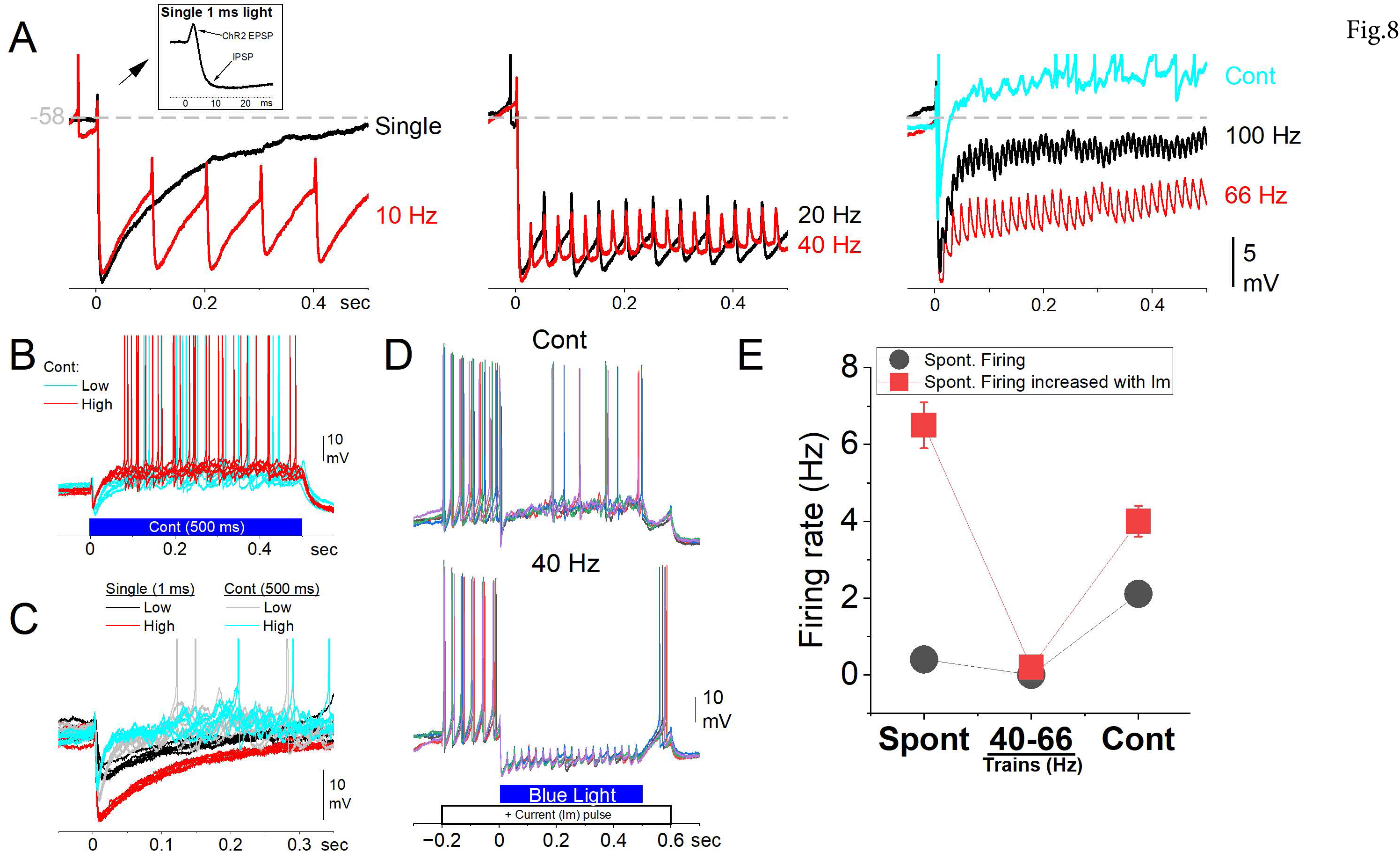
Optogenetic activation of ChR2-expressing NAc GABAergic neurons has complex effects on NAc neuron firing that depends on the light pattern. ***A,*** Whole-cell recordings from a NAc neuron in slices comparing the effects of different patterns of blue light on membrane potential (Vm). The black trace (single) and the inset on the left panel represent the effects of a single pulse of blue light revealing an EPSP followed by a sharp IPSP. Addition of repeated pulses in the form of trains at different frequencies augment and sustain the IPSPs. As the frequency of the trains increases, the EPSPs become more abundant and counteract the IPSPs. During Cont, the net effect is depolarizing, leading to action potential firing. ***B,*** Effect of Cont pulses of blue light at low and higher power. Cont pulses lead to NAc firing after the initial IPSP. ***C,*** Effect of single and Cont pulses at the low and higher light powers. Only the Cont pulses produce firing. ***D,*** Effect of Cont and 40 Hz trains on a NAc neuron activated by current injection. The trains completely inhibit the activated firing, while the Cont light supports firing after the initial IPSP, albeit at a lower level than the activated firing caused by current injection. **E,** Firing rate (Hz; n=3) comparing spontaneous (Spont) firing, 40-66 Hz trains, and Cont light during two conditions. Black circles represent cells firing without current injection and red squares represent the same cells activated by current injection. The trains robustly suppress firing regardless of activation level. Cont light increases cell firing when the neuron is not activated but inhibits firing when the neuron is activated albeit less efficiently than the trains.

These results indicate that trains and to some extent Cont blue light applied in the NAc of Vgat-NAc-ChR2 mice inhibits NAc neurons locally due to recurrent inhibition. Despite this inhibition of local somatodendritic NAc GABAergic neurons, the optogenetic light patterns should still directly stimulate the output axons of these neurons as they exit the NAc, just as they stimulate the synaptic collaterals causing local somatodendritic inhibition. If this is the case, the NAc GABAergic output axons should be activated by the blue light and inhibit distal targets outside of NAc. We proceeded to verify this next.

### Excitation of NAc GABAergic neurons inhibits NAc targets

Indirectly inhibiting a nucleus by activating local GABAergic neurons is straight-forward when the GABAergic neurons do not project outside the targeted nucleus. However, NAc GABAergic neurons project to various targets including ventral pallidum, SNr, and PPT (Gerfen and Surmeier, 2011; Castro and Bruchas, 2019). In a previous study (Hormigo et al., 2021c), we demonstrated that optogenetic excitation of NAc GABAergic axons in the SNr of Vgat-NAc-ChR2 mice inhibits both the SNr and PPT neurons they target. Here we tested if applying the blue light patterns within NAc has effects on distal targets in the midbrain due to the direct excitation of ChR2-expressing GABAergic axons exiting NAc, despite of the local NAc recurrent inhibition. If this is the case, the optogenetic light applied in the NAc of Vgat-NAc-ChR2 mice would directly inhibit the SNr, and consequently the superior colliculus (SC), which is under SNr tonic inhibitory control, would be disinhibited by the blue light applied in the NAc.

To investigate this hypothesis, we surgically implanted optical fibers in the NAc and a pair of movable microelectrodes (∼1 MΩ) in the SC of Vgat-NAc-ChR2 mice. Subsequently, we monitored the activity of SC cells while applying blue light patterns (Cont and trains at 66 Hz) in the NAc of freely moving mice.

Figure 9A,B plot the depth of the recording sites from the two electrodes (n=236 sites in 2 mice) versus the percent change in firing caused by 1 sec light pulses (combining all spikes from each site). Both Cont and trains of blue light applied in the NAc enhanced SC firing for cells located in a range of electrode depths that coincide with the SC intermediate and deep layers in histological sections along the electrode tracks referenced by marking lesions at known depths performed after the last recording session (Fig. 9A).

**Figure 9.**
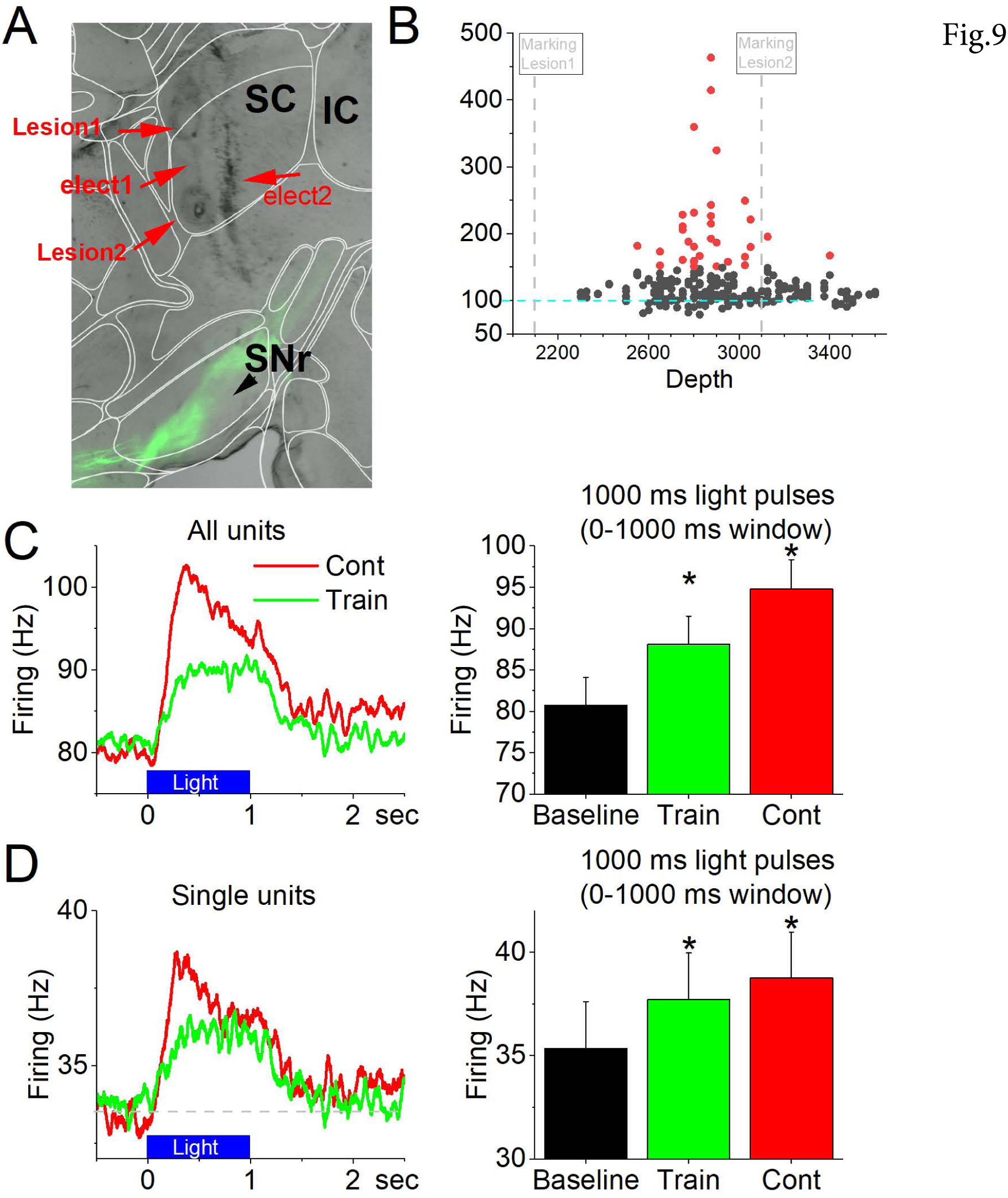
Optogenetic activation of ChR2-expressing NAc GABAergic neurons has downstream effects, such as SC disinhibition due to SNr cell inhibition. ***A***, Bright-field and fluorescent sagittal section aligned with the Allen Brain Atlas showing two marking lesions in the SC from one of the electrode pairs at a depth of 2000 and 3000 µm noted in ***B***. Note the fluorescent fibers in SNr originating from NAc GABAergic neurons, which also course toward the PPT area. ***B***, Effect of blue light Trains/Cont applied in the NAc of Vgat-NAc-ChR2 mice on the firing of SC cells per site (depth). The y-axis depicts the percent change in SC firing (all units combined per site) caused by 1 sec light pulses at the noted sites (depth). The results reveal a narrow range of depths where neuronal firing was affected by blue light applied in NAc. This narrow range coincides with the intermediate and deep layers of SC, which are under inhibitory control from SNr. The excitation of SC cells indicates that SNr cells are being inhibited by the blue light applied in the NAc. ***C***,***D***, PSTH of SC unit firing (Hz) for all (unsorted) units (***C***) or well-isolated single units (***D***) showing modulation caused by 1 sec Cont or Trains (40-66 Hz) of blue light applied in NAc. The 1 sec light Cont/light pulses recurred every 6 sec and were repeated at least 60 times per recording site. The population plots (right panels) measured firing during 1 sec windows prior to each light pulse (baseline), during the Cont and Train light. * p<0.05 vs baseline.

We then created PSTHs for all the units unsorted per electrode site (Fig. 9C; n=236), and for the sorted single units (Fig. 1D; n=246). There was an enhancement of SC firing during either Cont or Trains of blue light applied in the NAc for all the unsorted sites (Fig. 9C, Tukey t(234)> 9.4 p< 0.0001 Cont or Train vs Baseline) and the sorted single units (Fig. 9D, Tukey t(204)> 6.4 p< 0.0001 Cont or Train vs Baseline). The main difference between Cont and trains was the strong adaptation in firing caused by Cont which shows a stronger sharp enhancing effect at light onset that adapts rapidly. In contrast, trains enhanced firing less at onset but produced a sustained effect with little evidence of adaptation. These results indicate that the output axons of NAc neurons are being excited by both the trains and Cont blue light applied with the NAc of Vgat-NAc-ChR2 mice, which inhibit their targets (including SNr) causing further downstream effects. The enhanced NAc output will also inhibit PPT through direct fibers reaching this midbrain area, which we previously showed is sufficient to block signaled avoidance (Hormigo et al., 2019; Hormigo et al., 2021c).

### Excitation of NAc GABAergic neurons drives movement with an ipsiversive bias

In behaving rodents, asymmetric excitation of one side of the striatum, usually provoked by altering local dopamine, is associated with contraversive turning of the head/body (Leigh et al., 1983; Koshikawa, 1994). Contraversive turning also occurs during optogenetic activation of the direct output pathway from striatum (Kravitz et al., 2010). However, the effects of exciting the direct output pathway originating in NAc are less understood. In a recent paper, we found that unilateral excitation of NAc GABAergic ChR2-expressing axons in SNr produced no significant turning bias in either direction, while axons originating from dorsal and ventral portions of the striatum produced robust contraversive turning (Hormigo et al., 2021a; Hormigo et al., 2021c). Thus, the effects produced by striatum output fibers are not predictive of the effects of NAc output fibers.

Unilateral trains of blue light at 40-66 Hz applied in the NAc of Vgat-NAc-ChR2 mice (n=8) as they explored an open field produced an ipsiversive bias (Fig. 10A; Tukey t(6)= 7.17 p=0.002 40-66 Hz vs No Opsin). Unilateral Cont blue light also produced an ipsiversive bias but this was not significant compared to No Opsin mice (Fig. 10A; Tukey t(6)= 0.04 p=0.9), while 10-20 Hz trains (Tukey t(6)= 2.19 p=0.17) had little effect. Within the Vgat-NAc-ChR2 mice, both 40-66 Hz trains and Cont produced a stronger ipsiversive bias than 10-20 Hz trains (Tukey t(10)>3.89 p<0.05), and 40-66 Hz trains produced a stronger ipsiversive bias than Cont (Tukey t(10)=7.07 p=0.001).

**Figure 10.**
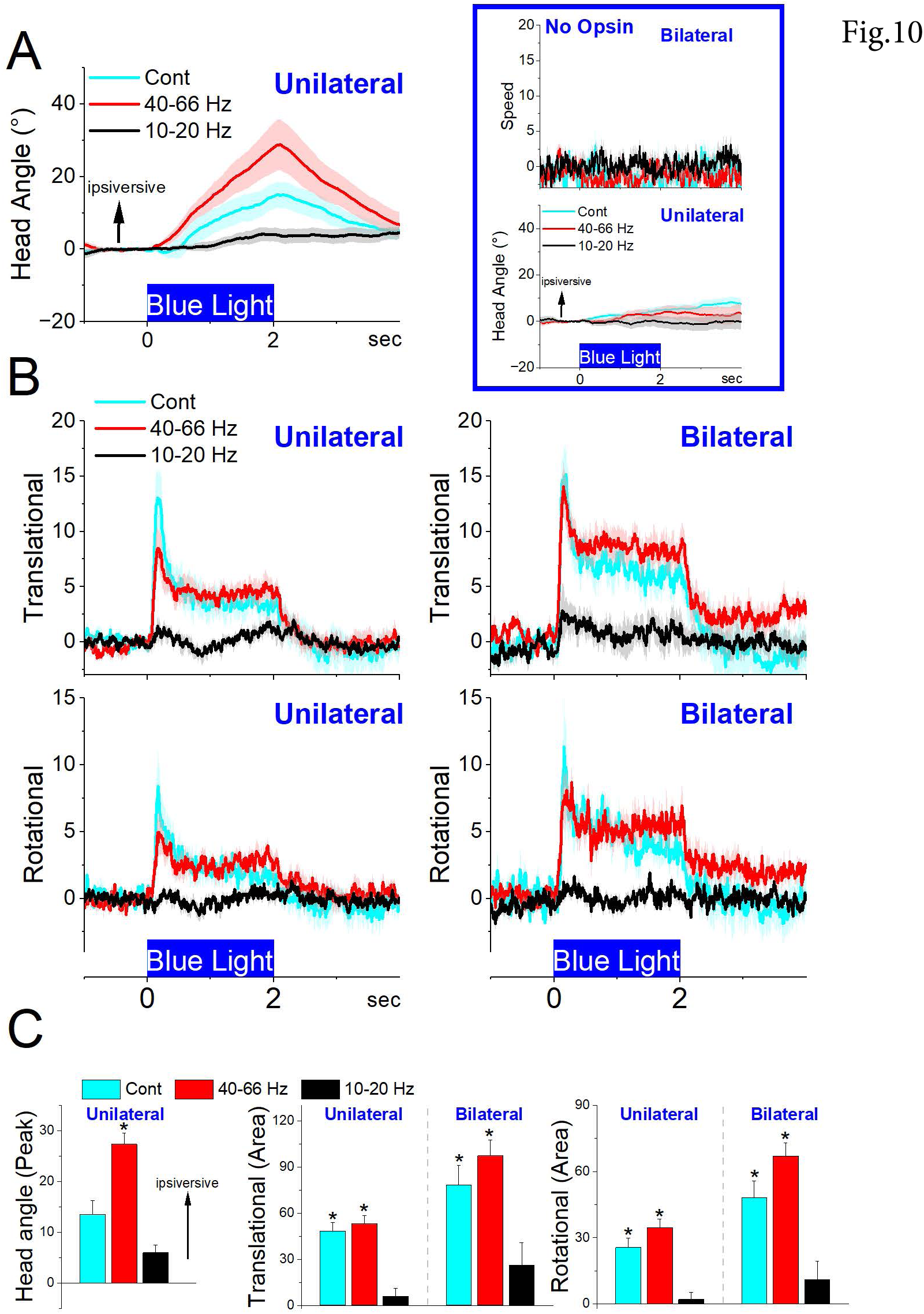
Optogenetic activation of ChR2-expressing NAc GABAergic neurons augments movement with an ipsiversive bias. ***A,*** Effect of blue light delivered unilaterally in the NAc of freely behaving Vgat-NAc-ChR2 mice in an open field to determine if the different patterns cause different turning/orienting biases (Angle). Low frequency trains (10-20 Hz) had little effect on orienting direction. In contrast, Cont and especially high frequency trains produced an ipsiversive bias. ***B,*** Same as ***A*** but shows the effect of light patterns on translational and rotational movement when the blue light is delivered unilaterally or bilaterally. During unilateral or bilateral light, both high frequency trains (40-66 Hz) and Cont increase translational and rotational movement. However, bilateral light produces more movement than unilateral light. ***C,*** Population measures for the results in ***A*** ang ***B***. * p<0.05 vs No Opsin mice subjected to the same light patterns.

We next measured the effect of unilateral and bilateral blue light trains on translational and rotational movement components without consideration of direction bias. Unilateral 10-20 Hz trains of blue light had little effect on translational and rotational movement. In contrast, unilateral 40-66 Hz trains of blue light generated translational (Tukey t(6)= 11.1 p=0.0002 40-66 Hz vs No Opsin) and rotational (Tukey t(6)= 10.5 p=0.0003) movement (Fig. 10B,C). Moreover, unilateral Cont blue light also generated translational (Tukey t(6)= 11.6 p=0.0001 Cont vs No Opsin) and rotational (Tukey t(6)= 8.2 p=0.001) movement. Within the Vgat-NAc-ChR2 mice, the amount of translational and rotational movement generated by 40-66 Hz trains and Cont blue light did not differ.

When 40-66 Hz trains or Cont light were applied bilaterally, both the translational (Tukey t(15)= 11.5 p<0.0001 40-66 Hz vs No opsin; Tukey t(15)= 7.08 p=0.0001 Cont vs No opsin) and rotational (Tukey t(15)= 10.4 p<0.0001 40-66 Hz vs No opsin; Tukey t(15)= 7.13 p=0.0001 Cont vs No opsin) movements increased robustly (Fig. 10B,C). Bilateral light increased translational movement compared to unilateral light during 40-66 Hz trains (Tukey t(40)= 5.9 p=0.004), but not during Cont (Tukey t(40)= 4.01 p=0.13) or 10-20 Hz trains. Bilateral light increased rotational movement compared to unilateral light during both 40-66 Hz trains (Tukey t(40)= 7 p=0.0004) and Cont (Tukey t(40)= 4.8 p=0.03). There was no difference in translational (Tukey t(40)= 2.47 p=0.71) or rotational (Tukey t(40)= 3.9 p=0.15) movement between 40-66 Hz trains and Cont bilateral light (Fig. 10B,C).

Taken together the results indicate that when blue light is applied unilaterally within NAc to activate NAc GABAergic neurons during exploration in an open field, movement increases with an ipsiversive bias. When the light is applied bilaterally, translational but especially rotational movement increases significantly. It is worth noting that bilateral blue light blocks active avoidance in these mice even though it generated more movement. Thus, the suppression of active avoidance cannot be explained by movement inhibition. However, the movement generated by bilateral light had a large rotational component, which may impair active avoidance because after the initial CS-driven orienting response, active avoidance responses consist primarily of translational movement, as mice shuttle in a forward motion between compartments (Zhou et al., 2023). The enhanced rotational component does not correspond to actual rotations but to exploratory-, orienting-like movements in random directions.

### Direct inhibition of NAc neurons has little effect on avoidance behaviors

The preceding results show that while optogenetic activation of NAc GABAergic neurons recurrently inhibits NAc locally, the output axons of these neurons are excited, with major effects downstream. Thus, we next determined the effect of directly optogenetically inhibiting NAc GABAergic neurons on signaled active avoidance.

To inhibit NAc GABAergic neurons, we expressed eArch3.0 in the NAc of Vgat-Cre mice with bilateral injections of a Cre-inducible AAV (Vgat-NAc-Arch, n=5 mice; Fig. 11A). In light trials, we tested the effects of 5 green light powers (AA1) or the higher light power (AA2, AA3). We found that, compared to CS trials, inhibiting NAc neurons in CS+Light trials had little effect on the percentage of active avoids as a function of green light power during AA1, AA2 and AA3-CS1 (Fig. 11A,B). Active avoids were unaffected by the powers tested during AA1 (Fig. 11A black symbols; RMAnova F(5,30)= 2.17 p=0.0841 CS vs CS+Light), AA2 (Fig. 11A red symbols; RMAnova F(1,8)= 1.03 p=0.3397), or AA3-CS1 (Fig. 11B left; Tukey t(27)= 1.61 p=0.66). Moreover, passive avoids were also unaffected during AA3-CS2 (Fig. 11B right; Tukey t(27)= 3.04 p=0.16).

**Figure 11.**
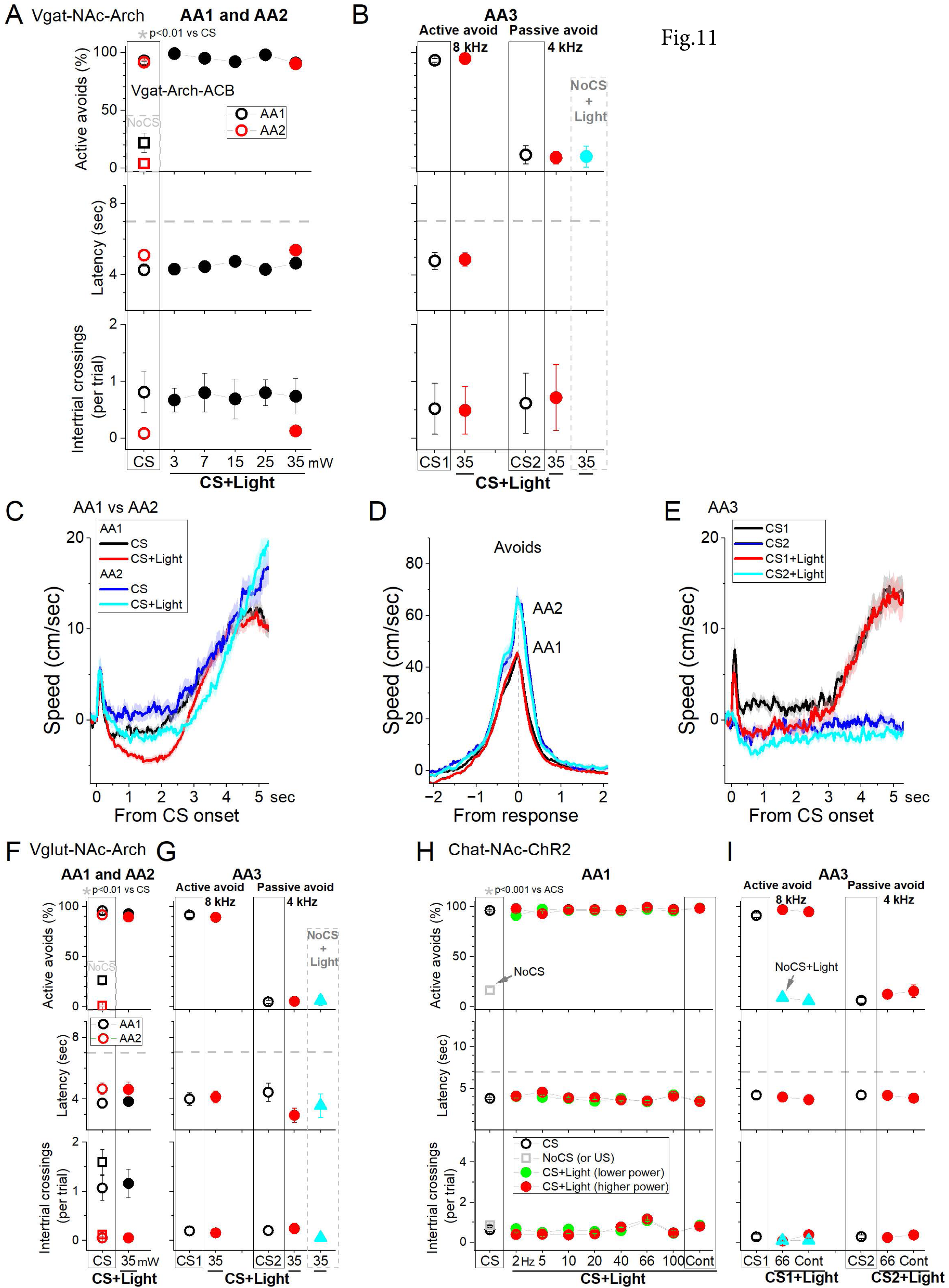
Optogenetic inhibition of NAc GABAergic neurons has little effect on signaled avoidance. ***A,B,*** Effect of Cont green light delivered at different powers on AA1/2 (***A***) and AA3 (***B***) in mice expressing eArch3.0 in NAc GABAergic neurons. ***C,*** Traces of overall movement (speed) during AA1 or AA2 for CS trials, and CS+Light trials combined. The trials are aligned by CS onset, which reveals the orienting response evoked by the CS followed by the ensuing avoid action. Note that CS+Light trials suppress ongoing movement prior to the onset of the avoid action without impairing the ability to respond. ***D,*** Same as ***C*** but traces are aligned by the avoidance response occurrence, which reveals the peak speed of the response. Note that avoids are not different between CS and CS+Light trials. As usual, AA2 produces faster avoids than AA1. ***E,*** Same as ***C*** but for AA3. CS1 and CS2 are control trials for active avoidance and passive avoidance, respectively. As per AA1 and AA2, ongoing movement is suppressed by the light. ***F,G,*** Same as ***A*** and ***B*** but for a control group of mice that do not express Arch in GABAergic NAc neurons as they were injected with an AAV targeting glutamatergic neurons. The light has negligible effects on AA1, AA2 (***F***), and AA3 (***G***). ***H,I,*** Effect of blue light on Chat-Cre mice that express ChR2 in NAc cholinergic neurons. Patterns of blue light were applied into NAc as in Fig. 7B,D during AA1 (***H***) and AA3(***I***). The light caused negligible effects on the performance of the mice.

Movement tracking revealed that inhibition of NAc GABAergic neurons had little effect on the orienting response during AA1, AA2 or AA3-CS2 trials (Fig. 11C,D,E; Tukey t(18)= 0.63 p=0.66). However, movement was suppressed after the orienting response and prior to the occurrence of the avoidance response (Fig. 11C; Tukey t(18)= 5.02 p=0.0023 CS vs CS+Light). Intriguingly, the ongoing movement pause caused by the light was not associated with impaired avoidance performance (Fig. 11A,B; avoids percentage and latency were unaffected). Thus, during the CS+Light trials mice pause right after the orienting response but avoid rapidly. There is no significant change in avoid peak speed measured from avoid occurrence (Fig. 11D; Tukey t(18)= 1.92 p=0.47 CS vs CS+Light from avoid occurrence).

As per the ChR2 experiments, we also tested control mice in which we injected the same Arch AAV in Vglut2 mice (n=2) or an Arch AAV with a CaMKII promoter in C57BL/6J mice (n=2) and combined these mice into a single control group (Fig. 11F,G, Glut-NAc-Arch, n=4 mice). In these mice, we found no effect of green light on active avoids during AA1 (Fig. 10F black symbols; RMAnova F(5,25)= 1.63 p=0.18), AA2 (Fig. 11F red symbols; RMAnova F(1,6)= 1.56 p=0.25), and AA3-CS1 (Fig. 11G left; Tukey t(28)= 2.55 p=0.39). We also found no effect on passive avoids during AA3-CS2 (Fig. 11G right; Tukey t(28)= 0.21 p=0.99) or NoCS+Light trials.

Since medium spiny GABAergic output neurons in NAc are controlled by cholinergic interneurons, which can inhibit NAc output when activated (de Rover et al., 2002; Castro and Bruchas, 2019), we expressed ChR2 by injecting an AAV in the NAc of Chat-Cre mice (Chat-NAc-ChR2, n=5). The results revealed little effect of exciting cholinergic NAc neurons with blue light on active avoids during AA1 (Fig. 11H, RMAnova F(16,48)= 1.28 p=0.2462), AA2 (RMAnova F(8,24)= 2.29 p=0.05), and AA3-CS1 (Fig. 11I, Tukey t(9)= 1.31 p=0.79), or on passive avoids during AA3-CS2 (Fig. 11I, Tukey t(9)= 2.49 p=0.34). Since these neurons are not known to project outside of NAc, the results indicate that the suppression of active avoidance caused in Vgat-NAc-ChR2 mice may be due to inhibitory effects outside of NAc.

Taken together, these results indicate that NAc neurons are not critically involved in signaled active or passive avoidance because directly inhibiting them has little effect on these behaviors.

### NAc lesions have little effect on avoidance behavior

To further determine if NAc neurons are important for learning and/or performing avoidance behaviors, we lesioned NAc GABAergic neurons by bilaterally injecting AAV8-EF1a-mCherry-flex-dtA into the NAc of Vgat-cre mice (n=9). To verify the lesion, we counted the number of neurons (Neurotrace) in the NAc lesion and control mice. The injection reduced the number of NAc neurons (Fig. 12A, Mann-Whitney Z=4.99 p<0.0001 Lesion vs Control). However, killing NAc GABAergic neurons had little effect on the ability of mice to learn and perform signaled active avoidance tasks (Fig. 12B). Mice learned AA1 at a rate like control mice. There was no difference in the percentage of active avoids between control and lesion mice during the AA1 (Anova F(1,13)= 0.03 p=0.85 Lesion vs Control), AA2 (F(1,13)= 0.02 p=0.89) or AA3-CS1 (F(1,13)= 0.53 p=0.48). There was also no difference in either avoidance latency (F(1,13)= 1.58 p=0.23) or intertrial crossings (F(1,13)= 0.03 p=0.85) between control and lesion mice during these active avoidance trials. Moreover, there was no difference in the percentage of passive avoids in AA3-CS2 (F(1,13)= 0 p=0.96) between control and lesion mice.

**Figure 12.**
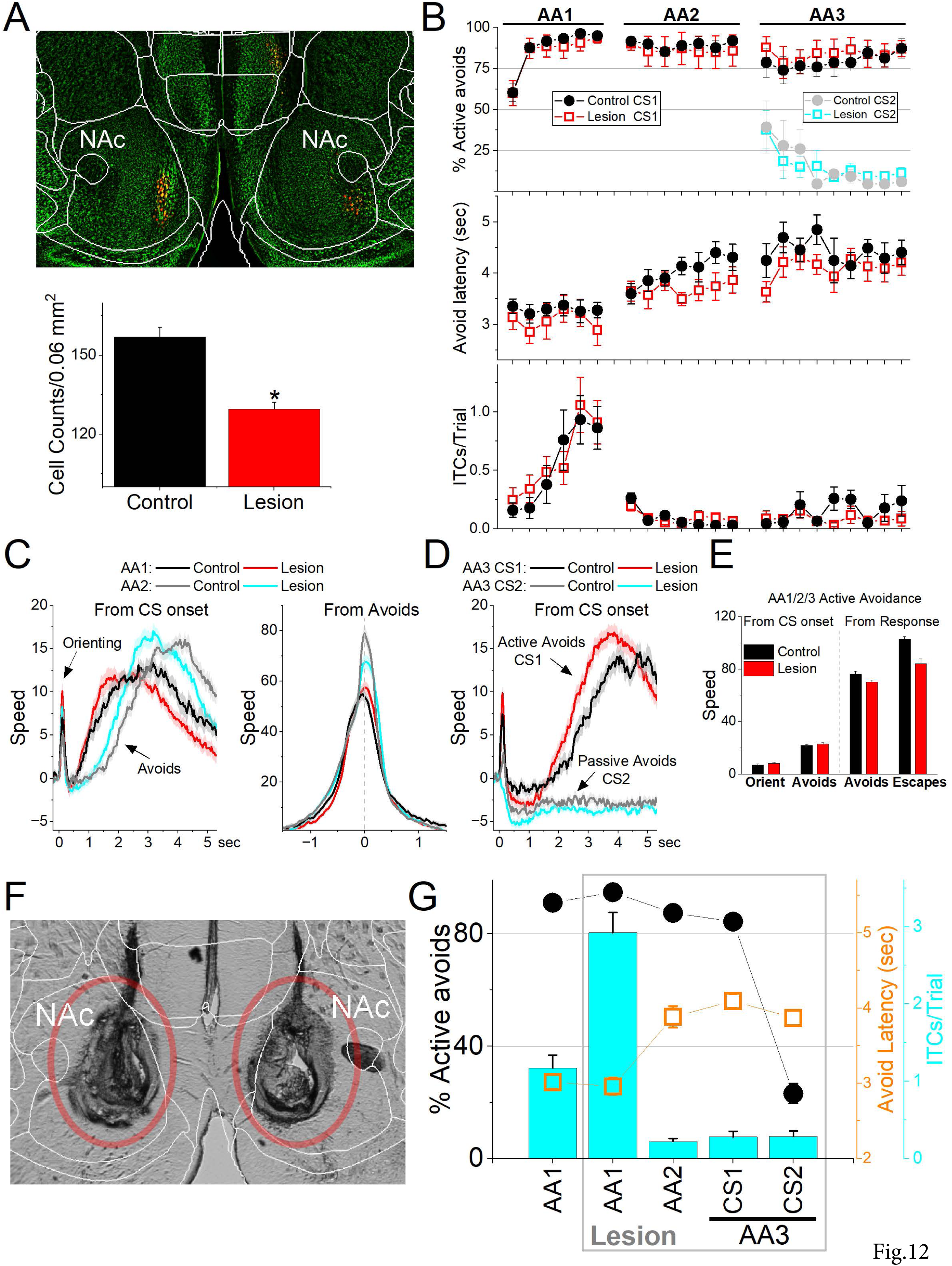
Lesions of NAc GABAergic neurons do not impair signaled avoidance learning or performance. ***A***, Coronal Neurotrace (green) stained section of a Vgat-Cre mouse injected with a Cre-dependent AAV-dtA in the NAc to kill GABAergic neurons. We counted the number of cells in the NAc in controls and lesion mice. There was a significant reduction in the number of NAc neurons in the lesion mice. ***B***, Behavioral performance during learning of AA1, followed by AA2 and AA3 procedures showing the percentage of active avoids (upper), avoid latency (middle), and ITCs (lower) for control and lesion mice. The AA3 procedure shows CS1 and CS2 trials for the same sessions. Active avoids during AA3-CS2 trials are errors, as the mice must passively avoid during CS2. Lesion mice tended to have shorter avoid latencies, but as is usual both control and lesion mice shifted their avoid latencies longer during AA2 compared to AA1. ***C***, Movement (speed) from CS onset (left) and from avoid occurrence (right) during AA1 and AA2 procedures for control and lesion mice. The lesion had little effect on movement other than the tendency to shift avoid movement sooner. However, the overall avoid speed measured from response occurrence was not faster in lesion mice. As typically observed in controls, lesion mice avoided faster during AA2 compared to AA1. ***D,*** Same as ***C*** for AA3. ***E***, Population measures of orienting, avoidance, and escape responses from CS onset (left) and from response occurrence (right) for overall movement. ***F***, Bilateral electrolytic lesions targeting the NAc. ***G***, Effect of bilateral electrolytic NAc lesions on behavioral performance in a repeated measures design. The plot shows the percentage of active avoids (filled black circles), avoid latency (open orange squares), and ITCs (cyan bars). Mice were trained in AA1 prior to the lesion and then placed back in AA1, followed by AA2 and AA3. The main effect of the lesion was to increase the number of ITCs during AA1, with little negative effect on active avoids. During AA2, lesion mice normally abolished their ITCs, which are punished, and shifted avoid latencies longer with little effect on avoidance performance. During AA3, lesion mice also performed AA3 normally, actively avoiding during CS1 and passively avoiding during CS2.

Comparison of movement (overall, rotational, and translational speed) during task performance did not show significant differences between lesion and control mice. Since these effects were similar in the AA1, AA2 and AA3 tasks, we combined the results for the three procedures (Fig. 12C,D,E). The orienting response evoked by the CS1 onset was not different between control and lesion mice (Fig. 12C,E; Anova F(1,13)= 0.06 p=0.80, including the rotational component F(1,13)= 0.12 p=0.73). The active avoidance movement speed was not different between control and lesion mice (Fig. 12C,E; F(1,13)= 0.4 p=0.53). This applied to both the translational (F(1,13)= 0.48 p=0.49) and rotational (F(1,13)= 0.01 p=0.90) movement. Similarly, when avoids failed, the escape response movement was not different between control and lesion mice (F(1,13)= 0.39 p=0.54). Finaly, the movement during passive avoids in AA3-CS2 trials did not differ between control and lesion mice (Fig. 12D; F(1,13)= 0.33 p=0.57). In conclusion, lesions of NAc GABAergic neurons had little effect on CS-evoked goal-directed movement driven by the CS and US.

Since the AAV-based lesion may leave some NAc cells intact, we performed electrolytic lesions of NAc (Fig. 12F), which assures elimination of NAc cells. We tested the effect of the lesion in trained mice (n=9; Fig. 12G) using a repeated measures design. The lesion did not affect the percentage of active avoids (F(1,8)= 0.61 p=0.45) or avoid latency (F(1,8)= 0.31 p=0.59) in AA1. However, there was a large increase in the number of ITCs (F(1,8)= 22.9 p=0.001), which can lead to spurious avoids. Subsequent training of mice in AA2, which abolishes ITCs and eliminates spurious avoids, revealed that mice continued to perform avoids at high rates (∼88%) without ITCs. Furthermore, the lesion mice learned AA3 like normal animals, which requires discriminating between CS1 and CS2 and selecting the appropriate action (Fig. 12G). These results indicate that NAc is not required to learn or perform active avoidance or to postpone action during passive avoidance. The lesions also had little impact on brain processes such as decision making (Go/NoGo in AA1-3), response inhibition (unsignaled and signaled passive avoidance in AA2/3), and stimulus discrimination (AA3).

## Discussion

We found that NAc neurons encode distinct facets of goal-directed movement during signaled avoidance. Calcium imaging showed that NAc GABAergic neurons activate during rotational exploratory movement, discerning the movement’s direction by exhibiting heightened discharges during contraversive head turning. Additionally, these neurons robustly discharge during signaled active avoidance behaviors, predominantly characterized by translational movement. Electrophysiological analysis confirmed the modulation of individual NAc neurons during CS-evoked orienting and active avoidance movement. These findings suggest a pivotal involvement of NAc neurons in signaled avoidance actions. Consequently, we employed optogenetics and lesions to manipulate the activity of NAc GABAergic neurons.

Activation of ChR2-expressing GABAergic NAc neurons resulted in a local interplay of excitation (direct EPSPs from ChR2 channels) and inhibition (indirect IPSPs from discharging ChR2-expressing GABAergic synapses), contingent on the pattern of applied blue light. Continuous blue light yielded early, adapting inhibition followed by net excitation, minimally impacting signaled avoidance. In contrast, high-frequency trains induced sustained NAc neuron recurrent inhibition, suppressing signaled avoidance while concurrently increasing overall movement. However, the indirect recurrent inhibition of NAc neurons by high-frequency trains and Cont is likely accompanied by the direct excitation of NAc output axons that target and inhibit NAc targets. Indeed, this was verified by showing downstream augmentation of neuronal firing in the midbrain SC caused by SNr inhibition upon light delivery in NAc. Thus, increased NAc GABAergic output inhibits NAc targets and suppresses signaled avoidance. This result is consistent with the effects of selectively exciting direct pathway axons originating in NAc that reach SNr or PPT in the midbrain, which also suppresses signaled avoidance (Hormigo et al., 2021c).

To determine the necessity of NAc during signaled avoidance, we directly inhibited NAc GABAergic neurons using Arch. Interestingly, this manipulation resulted in minimal impact on signaled avoidance despite a notable decrease in ongoing movement. Furthermore, electrolytic or AAV-based lesions of NAc demonstrated little effect on signaled avoidance. In summary, our findings indicate that while NAc neurons represent various facets of behavior related to movement, they do not play a central role in the generation of signaled avoidance. Nevertheless, under conditions of abnormal NAc functioning, leading to aberrant basal ganglia output, behavioral impairments may manifest, akin to observations in specific movement disorders.

### Complex effects of exciting networks of GABAergic neurons

We found that optogenetic excitation of ChR2-expressing NAc GABAergic neurons with high frequency trains of 1-ms blue light pulses recurrently inhibited NAc neurons by driving IPSPs. In contrast, Cont blue light pulses, excited quiescent NAc neurons or had less of an inhibitory effect than trains on active NAc neurons. The contrasting effects between trains and Cont blue light on networks of GABAergic neurons are consistent with previous findings. Optogenetic activation of networks comprising ChR2-expressing GABAergic neurons depends on the pattern of blue light applied (Hormigo et al., 2021a). Cont blue light directly induces depolarization and firing by persistently activating ChR2. In contrast, 40-66 Hz trains of 1-ms blue light pulses result in brief depolarizations that efficiently excite ChR2-expressing GABAergic synaptic fibers. This excitation, in turn, drives and sustains repeated IPSPs, leading to robust recurrent inhibition of cell firing (Hormigo et al., 2021a).

Despite the local recurrent inhibition within NAc, the blue light continues to excite NAc output fibers that inhibit NAc targets, such as ventral pallidum, SNr and PPT. Therefore, the inhibition of NAc GABAergic neurons with blue light trains in ChR2-expressing Vgat-NAc-ChR2 mice differs from the inhibition achieved with Arch in Vgat-NAc-Arch mice due to the activation of projection fibers beyond NAc in the Vgat-NAc-ChR2 mice.

Activation of NAc projection fibers led to increased movement with an ipsiversive bias, while direct inhibition of these NAc cells with Arch inhibited movement. The enhanced movement caused by increased NAc output likely results from activation of direct and indirect pathways (Kravitz et al., 2010; Gerfen and Surmeier, 2011; Castro and Bruchas, 2019). It may be mediated by NAc projections to the pallidum or other areas excluding SNr, because inhibition of SNr cells causes a contraversive movement bias in association with the disinhibition of SC (Hormigo et al., 2021a; Zhou et al., 2023).

### NAc neurons represent orienting and goal-directed avoidance movements

Taken together, the results indicate that NAc neurons are activated by movement, aligning with prior studies indicating that both direct and indirect striatal neurons typically remain silent in the absence of movement but activate during movement (Cui et al., 2013; Tecuapetla et al., 2014; Barbera et al., 2016). Striatal neurons continuously code movement variables, such as speed (Markowitz et al., 2018; Sales-Carbonell et al., 2018), likely driven by their sensitivity to somatosensory input (Carelli and West, 1991; Robbe, 2018).

NAc GABAergic neurons exhibited activation linked to exploratory movement, discerning orientation direction by discharging more strongly during contraversive turns. Notably, during auditory mapping sessions, these neurons discharged in response to auditory stimuli devoid of predictive behavioral contingencies. However, stimuli activating NAc neurons consistently coincided with movement events. This includes the orienting reflex, characterized by a rapid (<300 ms) head movement in response to salient auditory stimuli (Sokolov, 1963; Zhou et al., 2023). Beyond the swift onset activation of NAc neurons triggered by auditory tones, additional activation persisted throughout the 2-second duration, associated with pronounced exploratory rotational movements induced by the tones, particularly contraversive motions.

In the context of signaled avoidance, calcium imaging showed that NAc GABAergic neurons activate at CS onset in association with the orienting response, but then discharge more robustly during the ensuing active avoidance and escape movements. NAc activation has the potential to drive active avoids, but NAc activation may instead reflect the ongoing movement. We used several behavioral procedures to test the relationship between goal-directed behavior and NAc neuron activity. In general, NAc neurons were strongly activated during active avoidance movements, while passive avoids driven by auditory stimuli, which are not associated with movement, did not activate NAc neurons. Furthermore, unsignaled US presentations revealed a strong NAc activation during fast escape responses. Dissociation of the US components (footshock and auditory white noise) showed that the auditory stimulus adds little to the footshock evoked activation. Thus, NAc neurons activate in response to stimuli that produce movement regardless of the sensory modality driving the movement.

An interesting question is whether direct and indirect pathway NAc neurons activate distinctly during avoidance. One possibility is that direct pathway NAc neurons, traditionally associated with movement, would activate during active avoidance, while indirect pathway NAc neurons, traditionally associated with movement inhibition, would activate during passive avoidance. If this were the case, we would expect to see activation in NAc during passive avoidance reflecting the indirect pathway neurons. However, the only activation we observed was related to movement during active avoidance or other contexts, indicating that these cell groups may activate coherently during avoidance movement.

### NAc neurons are not required for goal-directed signaled avoidance

Direct inhibition of NAc GABAergic neurons with Arch in Vgat-NAc-Arch mice had little effects on signaled avoidance, although it significantly suppressed ongoing movement. The outcomes of direct inhibition in Vgat-NAc-Arch mice deviate from the anticipated effects that would be expected if the high-frequency blue light trains in Vgat-NAc-ChR2 mice were solely recurrently inhibiting the NAc. These two manipulations resulted in contrasting effects on signaled avoidance, orienting, and ongoing movement. This incongruity supports the notion that the high-frequency trains in Vgat-NAc-ChR2 mice do not solely recurrently inhibit NAc neurons locally, but also drive synchronous NAc output, leading to the dysregulation of NAc targets, which augments movement with an ipsiversive bias, and suppresses signaled avoidance. Furthermore, dysregulation of basal ganglia output is associated with movement disorders (Wichmann and Dostrovsky, 2011; Rubin et al., 2012; Willard et al., 2019). Moreover, activation of ChR2-expressing GABAergic Chat interneurons, which do not project outside of the NAc (Castro and Bruchas, 2019), had minimal impact on signaled avoidance like direct inhibition with Arch.

Lesions affecting the striatum, including the nucleus accumbens, can either enhance or impair active avoidance behaviors (Kirkby and Kimble, 1968; Darvas et al., 2011; Bravo-Rivera et al., 2014; Lichtenberg et al., 2014; Wendler et al., 2014; Ramirez et al., 2015). The striatum’s involvement in active avoidance has been recognized in both human and rodent studies (Oleson et al., 2012; Bravo-Rivera et al., 2015; Boeke et al., 2017). Our study agrees that NAc is engaged by active avoidance by coding aspects of the avoidance movement, including contraversive orienting. However, our study reveals that neither signaled active nor passive avoidance responses were affected by the direct inhibition or lesions of NAc GABAergic neurons. Although direct inhibition led to a marked suppression of ongoing movement, it did not impair the mice’s ability to execute active avoidance responses. Various goal-directed avoidance tasks involving distinct neural processes, including decision-making (Go/NoGo in AA1-3), action inhibition (unsignaled and signaled passive avoidance in AA2-3), and stimulus discrimination (AA3), were employed in our testing. The lesions or optogenetic inhibition demonstrated minimal impact on the mice’s proficiency in successfully performing these tasks.

Collectively, these findings suggest that NAc neurons do not play a critical role in either signaled active or passive avoidance behaviors. This aligns with the observation that the output of the basal ganglia through SNr is not inherently essential for mediating these behaviors under normal circumstances (Hormigo et al., 2021c). An important consideration is that while NAc is not requisite for normal avoidance behaviors, its dysregulation leading to aberrant output can significantly disrupt these behaviors by inhibiting its various targets, particularly in the midbrain. This may be relevant in the context of brain disorders, such as anxiety disorders leading to abnormal avoidance responses, and movement disorders disrupting signaled motions.

## Materials and Methods

### Experimental Design and Statistical Analysis

All procedures were reviewed and approved by the institutional animal care and use committee and conducted in adult (>8 weeks) male and female mice. Experiments involved a repeated measures design in which the mice or cells serve as their own controls (comparisons within groups), but we also compared experimental groups of mice or cells between groups (comparisons between groups). For comparisons within groups, we tested for a main effect of a variable (e.g., Stimulus) using a repeated measures ANOVA or a linear mixed-effects model with fixed-effects and random-effects (e.g., sessions nested within the Subjects as per Data ∼Stimulus + (1|Subjects/Sessions lme4 syntax in R) followed by comparisons with Tukey’s test. For comparisons between different groups, we used the same approach but included the Group as an additional fixed-effect (Data ∼Group*Stimulus + (1|Subjects/Sessions)). Using the standard errors derived from the model, Tukey tests were conducted for the effect of the fixed-effect factors (within group comparisons) or for the Group-Stimulus interaction (between group comparisons). We report the Tukey values for the relevant multiple comparisons. Plots report mean±SEM unless otherwise indicated.

To enable rigorous approaches, we maintain a centralized metadata system that logs all details about the experiments and is engaged for data analyses (Castro-Alamancos, 2022). Moreover, during daily behavioral sessions, computers run experiments automatically using preset parameters logged for reference during analysis. Analyses are performed using scripts that automate all aspects of data analysis from access to logged metadata and data files to population statistics and graph generation.

### Strains and Adeno-Associated Viruses (AAVs)

To record from NAc GABAergic neurons using calcium imaging, we injected a Cre-dependent AAV (AAV5-syn-FLEX-jGCaMP7f-WPRE (Addgene: 7×10^12^ vg/ml)) in the NAc of Vgat-cre mice (Jax 028862; B6J.129S6(FVB)-Slc32a1^tm2(cre)Lowl^/MwarJ) to express GCaMP6f/7f. An optical fiber was then placed in this location. To excite NAc GABAergic neurons using optogenetics, we employed Vgat-NAc-ChR2 mice in which we injected AAV5-EF1a-DIO-hChR2(H134R)-eYFP (UPenn Vector Core or Addgene, titers: 1.8×10^13^ GC/ml by quantitative PCR) in the NAc of Vgat-cre mice. To inhibit NAc GABAergic neurons using optogenetics, we expressed eArch3.0 by injecting AAV5-EF1a-DIO-eArch3.0-EYFP (UNC Vector Core, titers: 3.4×10^12^ vg/ml) in the NAc of Vgat-cre mice. In addition, to inhibit NAc neurons, we expressed ChR2 in cholinergic neurons of the NAc, which are GABAergic and inhibit principal GABAergic neurons locally, by injecting AAV5-EF1a-DIO-hChR2(H134R)-eYFP (UPenn Vector Core or Addgene, titers: 1.8×10^13^ GC/ml) into the NAc of Chat-cre mice (Jax 031661; B6.129S-Chat^tm1(cre)Lowl^/MwarJ). No-Opsin controls were injected with AAV8-hSyn-EGFP (Addgene, titers: 4.3×10^12^ GC/ml by quantitative PCR) or nil in the NAc.

As additional controls of the optogenetic effects of manipulating the activity of GABAergic (Vgat) NAc neurons, we targeted Vglut2 or CaMKII in NAc. For Arch controls, we injected AAV5-EF1a-DIO-eArch3.0-EYFP (UNC Vector Core, titers: 3.4×10^12^ vg/ml) in the NAc of Vglut2-cre mice and injected AAV5-CaMKIIa-eArchT3.0-EYFP (UNC Vector Core, titers: 4×10^12^ vg/ml) in the NAc of C57BL/6J mice. For ChR2 controls, we injected AAV5-EF1a-DIO-hChR2(H134R)-eYFP (UPenn Vector Core or Addgene, titers: 1.8×10^13^ GC/ml by quantitative PCR) in the NAc of Vglut2-cre mice and injected AAV5-CaMKIIa-hChR2(H134R)-EYFP (UNC Vector Core, titers: 6.2×10^12^ vg/ml) in the NAc of C57BL/6J mice. In accordance with the lack of glutamatergic (e.g., Vglut2) cells in NAc, these controls resulted in nil expression in the NAc (Vong et al., 2011).

For optogenetics, we implanted dual optical fibers bilaterally in the NAc. All the AAVs and optogenetic methods used in the present study have been validated in previous studies using slice and/or in vivo electrophysiology (Hormigo et al., 2016; Hormigo et al., 2019; Hormigo et al., 2021a; Hormigo et al., 2021c).

### Surgeries

Optogenetics and fiber photometry experiments involved injecting 0.2-0.4 µl AAVs per site during isoflurane anesthesia (∼1%). Animals received carprofen after surgery. The stereotaxic coordinates for injection in NAc are (in mm from bregma; lateral from the midline; ventral from the bregma-lambda plane): 1.2 anterior, 0.8, and 4.3. In these experiments, a single (400 µm in diameter for fiber photometry) or dual (200 µm in diameter for optogenetics) optical fiber was implanted unilaterally or bilaterally during isoflurane anesthesia. The stereotaxic coordinates for the implanted optical fibers (in mm) in the NAc are: 1.2 anterior, 1, and 4.3. No Opsin mice were implanted with cannulas in NAc or adjacent sites.

Bilateral electrolytic lesions were performed by lowering an electrode (0.25 mm diameter, insulated stainless steel with a 0.5 mm exposed tip) into the NAc and applying current (0.6 mA) for 15 sec in an anterior (1.0 anterior, 0.8 lateral, and 4.6 ventral in mm) and a posterior site (1.6 anterior, 0.8, and 4.6 mm) per side.

In unit recording experiments, a bundle of several electrodes including a ground, analog reference, and tungsten sharp (edged) electrodes (1-2 MΩ) was implanted in the NAc or SC. The ground and analog reference were placed outside the NAc or SC, and a blunt tungsten electrode served as a digital reference located near the recording electrodes. Implanted recording electrodes were static (NAc) or movable (SC) using an implanted screw-based drive manipulator.

### Active Avoidance tasks

Mice were trained in a signaled active avoidance task, as previously described (Hormigo et al., 2016; Hormigo et al., 2019). During an active avoidance session, mice are placed in a standard shuttle box (16.1” x 6.5”) that has two compartments separated by a partition with side walls forming a doorway that the animal must traverse to shuttle between compartments. A speaker is placed on one side, but the sound fills the whole box and there is no difference in behavioral performance (signal detection and response) between sides. A trial consists of a 7 sec avoidance interval followed by a 10 sec escape interval. During the avoidance interval, an auditory CS (8 kHz, 85 dB) is presented for the duration of the interval or until the animal produces a conditioned response (avoidance response) by moving to the adjacent compartment, whichever occurs first. If the animal avoids, by moving to the next compartment, the CS ends, the escape interval is not presented, and the trial terminates. However, if the animal does not avoid, the escape interval ensues by presenting white noise and a mild scrambled electric foot-shock (0.3 mA) delivered through the grid floor of the occupied half of the shuttle box. This unconditioned stimulus (US) readily drives the animal to move to the adjacent compartment (escape response), at which point the US terminates, and the escape interval and the trial ends. Thus, an *avoidance response* will eliminate the imminent presentation of a harmful stimulus. An *escape response* is driven by presentation of the harmful stimulus to eliminate the harm it causes. Successful avoidance warrants the absence of harm. Each trial is followed by an intertrial interval (duration is randomly distributed; 25-45 sec range), during which the animal awaits the next trial. We employed four variations of the basic signaled active avoidance procedure termed AA1, AA2, AA3 and AA4.

In AA1, mice are free to cross between compartments during the intertrial interval; there is no consequence for intertrial crossings (ITCs).

In AA2, mice receive a 0.2 sec foot-shock (0.3 mA) and white noise for each ITC. Therefore, in AA2, mice must passively avoid during the intertrial interval by inhibiting their tendency to shuttle between trials. Thus, during AA2, mice perform both signaled active avoidance during the signaled avoidance interval (like in AA1) and unsignaled passive avoidance during the unsignaled intertrial interval.

In AA3, mice are subjected to a CS discrimination procedure in which they must respond differently to a CS1 (8 kHz tone at 85 dB) and a CS2 (4 kHz tone at 70 dB) presented randomly (half of the trials are CS1). Mice perform the basic signaled active avoidance to CS1 (like in AA1 and AA2), but also perform signaled passive avoidance to CS2, and ITCs are not punished. In AA3, if mice shuttle during the CS2 avoidance interval (7 sec), they receive a 0.5 sec foot-shock (0.3 mA) with white noise and the trial ends. If animals do not shuttle during the CS2 avoidance interval, the CS2 trial terminates at the end of the avoidance interval (i.e., successful signaled passive avoidance).

In AA4, three different CS’s, CS1 (8 kHz tone at 81 dB), CS2 (10 kHz tone at 82 dB), and CS3 (12 kHz tone at 82 dB) signal a different avoidance interval duration of 4, 7, and 15 sec, respectively. Like in AA2, mice are punished for producing intertrial crossings. In AA4, mice adjust their response latencies according to the duration of the avoidance interval signaled by each CS.

There are three main variables representing task performance. The percentage of active avoidance responses (% avoids) represents the trials in which the animal actively avoided the US in response to the CS. The response latency (latency) represents the time (sec) at which the animal enters the safe compartment after the CS onset; avoidance latency is the response latency only for successful active avoidance trials (excluding escape trials). The number of crossings during the intertrial interval (ITCs) represents random shuttling due to locomotor activity in the AA1 and AA3 procedures, or failures to passively avoid in the AA2 procedure. The sound pressure level (SPL) of the auditory CS’s were measured using a microphone (PCB Piezotronics 377C01) and amplifier (x100) connected to a custom LabVIEW application that samples the stimulus within the shuttle cage as the microphone rotates driven by an actuator controlled by the application.

### Unit recordings

Mice were connected to a rotatory-assisted electrical swivel by an ultra-light cable and digital headstage (<1 g) and were free to move in the shuttle box. Activity from all the implanted electrodes was continuously recorded as wide-band, high-pass and low-pass filtered signals. Unit activity was threshold detected and sorted using PCA, and k-means clustering (*scikit-learn*) manually or with automated scripts (validated with manual sorting). For movable electrodes, the depth was adjusted prior to each recording session to find a well-isolated unit within the target area. After the last recording session, marking lesions (DC, 10 sec 0.1 mA) were made at the static electrode locations or at defined depths of the movable electrodes for histological verification.

### Fiber photometry

We employed a 2-channel excitation (465 and 405 nm) and 2-channel emission (525 and 430 nm for GCaMP6f and control emissions) fiber photometry system (Doric Lenses). Alternating light pulses were delivered at 100 Hz (per each 10 ms, 465 is on for 3 ms, and 2 ms later 405 is on for 3 ms). While monitoring the 525 nm emission channel, we set the 465 light pulses in the 20-60 µW power range and then the power of the 405 light pulses was adjusted (20-50 µW) to approximately match the response evoked by the 465 pulses. During recordings, the emission peak signals evoked by the 465 (GCaMP6f) and 405 (isobestic) light pulses were acquired at 5-20 kHz and measured at the end of each pulse. To calculate F_o_, the measured peak emissions evoked by the 405 nm pulses were scaled to those evoked by the 465 pulses (F) using the slope of the linear fit. Finally, F/F_o_ was calculated with the following formula: (F-F_o_)/F_o_ and converted to Z-scores. Due to the nature of the behavior studied, a swivel is essential. We employed a rotatory-assisted photometry system that has no light path interruptions (Doric Lenses). In addition, black acrylic powder was mixed in the dental cement to assure that ambient light was not leaking into the implant and reaching the optical fiber; this was tested in each animal by comparing fluorescence signals in the dark versus normal cage illumination.

### Optogenetics

The implanted optical fibers were connected to patch cables using sleeves. A black aluminum cap covered the head implant and completely blocked any light exiting at the ferrule’s junction. Furthermore, the experiments occurred in a brightly lit cage that made it difficult to detect any light escaping the implant. The other end of the patch cable was connected to a dual light swivel (Doric lenses) that was coupled to a green laser (520 nm; 100 mW) to activate Arch or a blue laser (450 nm; 80 mW) to activate ChR2. In experiments expressing Arch, Green light was applied continuously at different powers (3.5, 7, 15, 25 and 35 mW). In experiments expressing ChR2, blue light was applied at the same power in different patterns, including continuous (Cont) and in trains of 1-ms pulses at different frequencies (2-100 Hz). The blue light included low (∼1 mW), medium (∼3 mW), and high (∼6 mW) powers. Unless otherwise noted, if the effects of low and medium blue light power were similar, the results were combined. Power is regularly measured by flashing the connecting patch cords onto a light sensor – with the sleeve on the ferrule.

During optogenetic experiments that involve avoidance procedures, we compared five different trial types: CS, CS+Light, Light, NoCS, and NoCS+Light trials. *CS trials* were normal trials for a particular procedure without optogenetic light. *CS+Light trials* were identical to CS trials except that optogenetic light was delivered simultaneously with the CS and US (if delivered) during the avoid and escape intervals, respectively. *Light trials* were identical to CS+Light trials but the CS was omitted to determine if the Light could serve the same signaling function as the CS. *NoCS trials* were catch trials that lack CS or US to measure responses delivered by chance. *NoCS+Light trials* are identical to NoCS trials, but light was delivered to determine its sole effects. To perform within group repeated measures (RM) comparisons, the different trial types for a procedure were delivered randomly within the same session. In addition, the trials were compared between different groups, including No Opsin mice that did not express opsins but were subjected to the same trials including light delivery.

### Video tracking

During open field experiments, mice are placed in a circular open field (10” diameter) that was illuminated from the bottom or in a standard shuttle box (16.1” x 6.5”). All mice in the study (open field or shuttle box) were continuously video tracked (30-100 FPS) in synchrony with the procedures and other measures. We automatically tracked head movements with two color markers attached to the head connector –one located over the nose and the other between the ears. The coordinates from these markers form a line (head midline) that serves to derive several instantaneous movement measures per frame (Zhou et al., 2023). Overall head movement was separated into *rotational* and *translational* components (unless otherwise indicated, overall head movement is presented for simplicity and brevity, but the different components were analyzed). Rotational movement was the angle formed by the head midline between succeeding video frames multiplied by the radius. Translational movement resulted from the sum of linear (forward vs backward) and sideways movements. *Linear* movement was the distance moved by the ears marker between succeeding frames multiplied by the cosine of the angle formed by the line between these succeeding ear points and the head midline. *Sideways* movement was calculated as linear movement, but the sine was used instead of the cosine. Pixel measures were converted to metric units using calibration and expressed as speed (cm/sec). We used the time series to extract window measurements around events (e.g., CS presentations). Measurements were obtained from single trial traces and/or from traces averaged over a session. In addition, we obtained the direction of the rotational movement with a *Head Angle* or *bias* measure, which was the accumulated change in angle of the head per frame (versus the previous frame) zeroed by the frame preceding the stimulus onset or event (this is equivalent to the rotational speed movement in degrees). The *time to peak* is when the *extrema* occurs versus event onset.

To detect spontaneous turns or movements from the head tracking, we applied a local maximum algorithm to the continuous head angle or speed measure, respectively. Every point is checked to determine if it is the maximum or minimum among the points in a range of 0.5 sec before and after the point. A change in angle of this point >10 degrees was a detected turn in the direction of the sign. We further sorted detected turns or movements based on the timing of previous detected events.

### Histology

Mice were deeply anesthetized with an overdose of ketamine or isoflurane. Upon losing all responsiveness to a strong tail pinch, the animals were decapitated, and the brains were rapidly extracted and placed in fixative. The brains were sectioned (100 µm sections) in the coronal or sagittal planes. Some sections were stained using Neuro-trace or cresyl violet. All sections were mounted on slides, cover slipped with DAPI mounting media, and all the sections were imaged using a slide scanner (Leica Thunder). We used an APP we developed with OriginLab (Brain Atlas Analyzer) to align the sections with the Allen Brain Atlas Common Coordinate Framework (CCF) v3 (Wang et al., 2020). This reveals the location of probes, fluorophores, and lesions versus the delimited atlas areas. We used it to delimit Neurotrace or cresyl violet stained sections of electrolytic lesion mice and determine the extent of the lesions.

## Acknowledgments

Supported by NIH grants to MAC. We thank Mariana Mangini for technical assistance. Additional information at castro-lab.org.

